# Arginine methylation sites on SepIVA help balance elongation and septation of the cell wall in *Mycobacterium smegmatis*

**DOI:** 10.1101/2021.10.06.463415

**Authors:** Angela H Freeman, Karen Tembiwa, James R Brenner, Michael R Chase, Sarah M Fortune, Yasu S Morita, Cara C Boutte

## Abstract

Growth of mycobacterial cells requires successful coordination between elongation and septation of the cell wall. However, it is not clear which factors directly mediate this coordination. Here, we studied the function and post-translational modification of an essential division factor, SepIVA, in *Mycobacterium smegmatis*. We find that SepIVA is arginine methylated, and that alteration of these methylation sites affects both septation and polar elongation of *Msmeg*. Furthermore, we show that SepIVA regulates the localization of MurG, and that this regulation may impact polar elongation. Finally, we map SepIVA’s two regulatory functions to different sites on the protein: the N-terminus regulates elongation while the C-terminus regulates division. These results establish SepIVA as a regulator of both elongation and division and characterize a physiological role for protein arginine methylation sites for the first time in mycobacteria.

## Introduction

The cell cycle of rod-shaped bacteria requires coordination of elongation and division. Elongation involves incorporation of newly synthesized cell wall components, including peptidoglycan (PG) into the existing cell wall. In the well-studied species *B. subtilis* and *E. coli*, this incorporation occurs along the lateral cell walls and is directed by the cytoskeletal factor MreB (Egan *et al*., 2020). However, some rod-shaped bacteria, including many Actinobacteria and some Alpha-proteobacteria, lack MreB-like proteins and elongate by incorporating new cell wall near the cell poles (Thanky *et al*., 2007; Aldridge *et al*., 2012; Brown *et al*., 2012; Cameron *et al*., 2015; Zupan *et al*., 2019; Zupan *et al*., 2021). Polar elongation in mycobacteria, which includes the pathogen *Mycobacterium tuberculosis*, requires the DivIVA protein Wag31 (Kang, 2005; Kang *et al*., 2008; Melzer *et al*., 2018), through mechanisms that are currently unclear.

Many essential cell division factors are conserved from *E. coli* to mycobacteria (Hett and Rubin, 2008; Wu *et al*., 2018). However, because of the extra layers of the cell wall (Jankute *et al*., 2015), and polar growth (Thanky *et al*., 2007) in mycobacteria, we expect that some aspects of cell division must be different. In mycobacteria, polar elongation and septation are spatially coincident but temporally separated; while in lateral growers, elongation and septation are spatially distinct. Because division creates the site of polar growth for the next generation of mycobacterial cells, we expect there to be proteins involved in transitioning the new pole from the septal/divisive mode cell wall metabolism (orthogonal to the long axis of the cell) to the elongative mode of cell wall metabolism (parallel to the long axis of the cell) - though such factors have not been described.

The intracellular membrane domain (IMD) is likely shared by the elongation and division processes. The IMD is a membranous hub for generation of lipid-linked cell wall precursors and it localizes to sites of cell wall synthesis: the subpolar region and septum (Hayashi *et al*., 2016). Cell wall precursors assembled in the IMD are thought to be translocated into the plasma membrane associated with the cell wall (PM-CW) before assembly into the cell wall (García-Heredia *et al*., 2021). How precursor enzymes associate with the IMD, and how that affects their activity is unknown.

SepIVA is essential for division and interacts with the conserved divisome protein, FtsQ (Wu *et al*., 2018; Jain *et al*., 2018). *sepIVA* has also been implicated as a determinant of beta-lactam susceptibility, indicating that it may affect peptidoglycan metabolism (Flores *et al*., 2005). Like Wag31, SepIVA comprises a DivIVA domain and aligns to DivIVA from *B. subtillis* (Fig. S1). DivIVA homologs are often involved in recruitment of other proteins to cell poles or septa (Marston and Errington, 1999; Halbedel and Lewis, 2019; Hammond *et al*., 2019; Hammond *et al*., 2021). However, none of the confirmed interactors of DivIVA-domain proteins interact with residues that are conserved in SepIVA (Fig. S1). Depletion of Wag31 and SepIVA have opposite effects: depletion of SepIVA leads to continued elongation and defective division (Wu *et al*., 2018), while depletion of Wag31 leads to elongation arrest but continued division (Kang *et al*., 2008).

Here, we show that SepIVA is arginine methylated. Arginine methylation is known to be involved in various cell functions in eukaryotes, such as membrane association, protein transport, DNA binding and protein-protein interactions (Biggar and Li, 2015; Raposo and Piller, 2018). Protein arginine methylation has been identified using mass spectrometry in *E. coli* (Zhang *et al*., 2018), mycobacteria (Sakatos *et al*., 2018) and *Salmonella* (Su *et al*., 2021). Arginine methylation sites have been shown *in vitro* to affect DNA binding of transcription factors in *Salmonella* (Su *et al*., 2021) and mycobacteria (Singhal *et al*., 2020). However, the physiological roles of protein arginine methylation have barely been studied in bacteria. Currently, there are no characterized protein arginine methyltransferases in mycobacteria. Recent work has suggested that the protein glutamate methyltransferase CheR in *Salmonella enterica* can somehow also methylate arginines (Su *et al*., 2021); there is no homolog with a predicted CheR methyltransferase domain in mycobacteria.

In this work, we show that methyl-ablative and methyl-mimetic mutations at certain arginine methylation sites on SepIVA regulate SepIVA’s localization, peptidoglycan metabolism, and cell elongation and division. We also show that SepIVA is involved in regulating the localization of peptidoglycan precursor enzyme MurG, and that arginine methylation site mutations of *sepIVA* affect MurG’s localization to the poles. This work establishes SepIVA as a regulator of both elongation and division and indicates that it may have a role in regulating MurG. In addition, we show here that SepIVA is arginine methylated, and that methyl-mimetic mutations at the N-terminus of SepIVA disrupt MurG’s polar localization and slow polar growth, while methyl-ablative site mutations promote polar growth. We also find that methylation sites near the C-terminus of SepIVA are required for normal cell division.

## Results

### Methylation site mutations on SepIVA affect growth and cell length of Msmeg

Post-translational modifications are critical for regulating the physiology of *Mtb* and other mycobacteria (Kang *et al*., 2008; Prisic *et al*., 2010; Garces *et al*., 2010; Prisic and Husson, 2014; Boutte *et al*., 2016; Sakatos *et al*., 2018; Iswahyudi *et al*., 2019; Shamma *et al*., 2020). Most studies thus far have focused on phosphorylation. To identify proteins that could be regulated by other types of post-translational modification, we re-analyzed a mass spectrometry data set of *Mtb* peptide masses that we had previously published (Garces *et al*., 2010) to search for post translational modifications. We found evidence of four arginine methylations on SepIVA (Rv2927c), as well as on many other proteins from *Mtb* (Supplemental table 1). The four methylated arginines from *Mtb* SepIVA were R19, R105, R111, R199 and all are conserved in SepIVA from *Msmeg*.

To see if SepIVA in *M. smegmatis* is also arginine methylated, we immunoprecipitated SepIVA-strep from *Msmeg* lysates and used LCMS-IT-TOF mass spectrometry to measure the post-translational modifications. We found evidence of widespread arginine methylation on SepIVA’s 24 arginine residues (Supplemental table 2).

To identify arginine residues in SepIVA for which methylation might have a physiological role, we used L5 allele swapping to make several mutants of *sepIVA* in *Msmeg* (Pashley and Parish, 2003). First, we integrated a wild-type *sepIVA* at the L5 site on a nourseothricin-marked vector and deleted the endogenous copy of *sepIVA*. Then, we cloned various *sepIVA* arginine mutants into kanamycin-marked vectors, and transformed these into the Δ*sepIVA L5::sepIVA* WT strain, and selected for allele swaps. Because there were so many arginines with potential methylation sites, we made mutants with several clustered arginines mutated at once. We grouped arginines based on their location on the SepIVA protein, as predicted by AlphaFold (Jumper *et al*., 2021) (Pettersen *et al*., 2021) (Fig. S2). We made R->M methionine mutants to represent methyl-mimetic mutations, as methionine is the closest structural mimic to a methylated arginine (Singhal *et al*., 2020). We made R->A alanine mutations to test the function of the positive charge of arginines and R->K lysine mutations to serve as methyl-ablatives that retain the positive charge on the residue. We found that several groups of arginines were essential for *sepIVA* function, as the alanine mutant strains did not grow (R->A mutants, Fig. 1A) in liquid culture.

**Figure 1.**
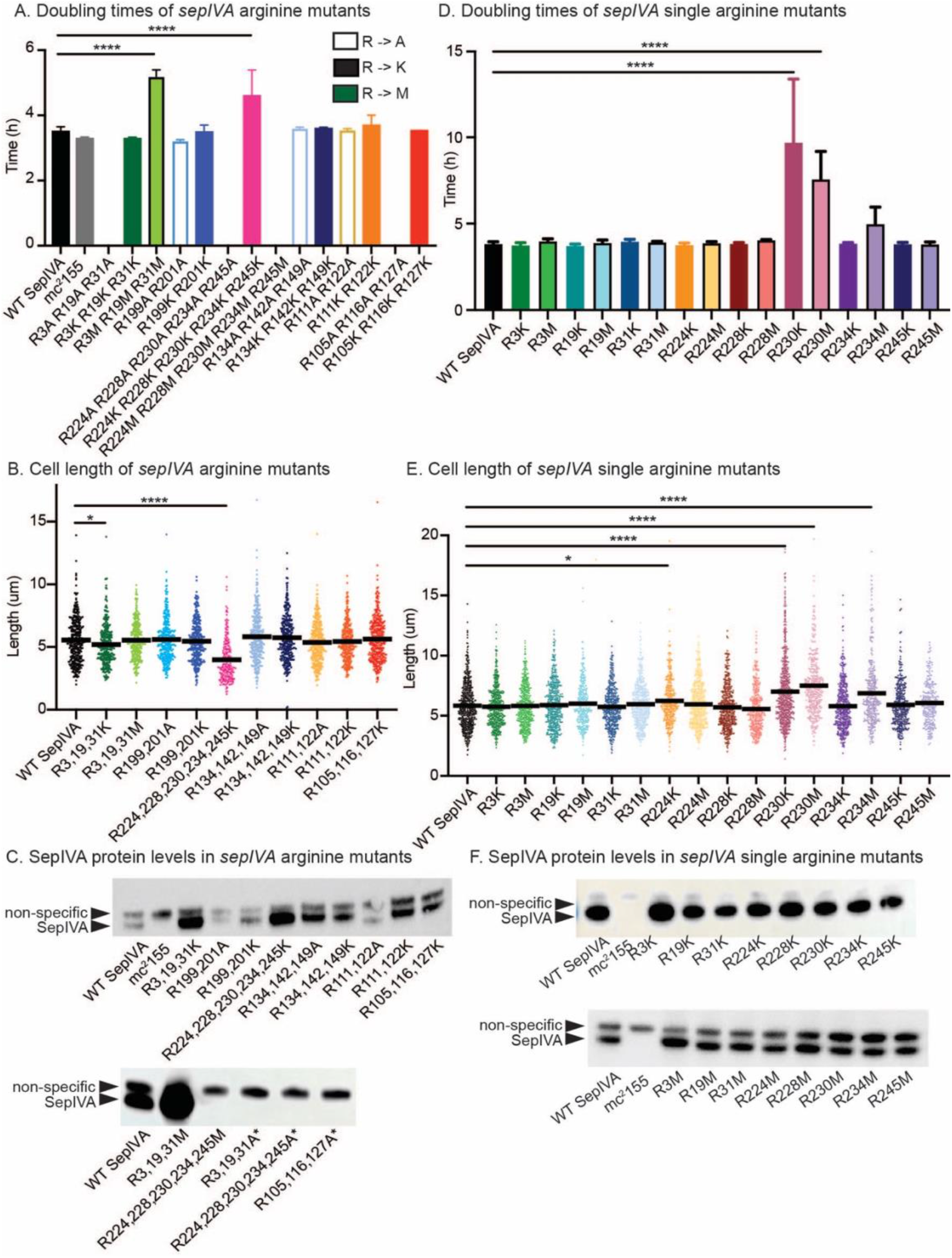
Methylation site mutations on SepIVA affect growth and cell length of *Msmeg*. **(A)** Doubling time of allele-swap *M. smeg* cells expressing WT or arginine methylation site mutants of SepIVA. WT SepIVA represents the L5 complemented strain. Lighter color outlined bars represent arginine to alanine mutants. No outline, darker color bars represent methyl-ablative methylation site mutants. Black outline bars represent methyl-mimetic methylation site mutants. Strains without bars were non-viable. At least 3 biological replicates of each strain were used. Error bars represent standard deviation. ****, *P = <0.0001*. **(B)** Cell lengths of SepIVA WT or arginine methylation site mutants. Bars represent mean. At least 300 cells over three biological replicates of each strain were analyzed. *, *P = 0.0439; ****, P = <0.0001*. **(C)** α-strep western blot of L5::*sepIVA-*strep WT or arginine methylation site mutants. mc^2^155 without a strep tag was used as a negative control (lane 2). The non-specific band serves as a loading control. ‘*’ denotes merodiploid strains. Merodiploid strains were used for SepIVA variants did not support growth as allele swaps. **(D)** Doubling time of allele-swap *M. smeg* cells expressing WT or single arginine methylation site mutants of SepIVA. WT SepIVA is the strain with WT *sepIVA*-strep at the L5 site and *sepIVA* deleted from its native locus – this is the isogenic control. No outline, darker color bars represent methyl-ablative methylation site mutants. Black outline bars represent methyl-mimetic methylation site mutants. Strains without bars were non-viable. At least 3 biological replicates of each strain were used. Error bars represent standard deviation. ****, *P = <0.0001*. **(E)** Cell lengths of SepIVA WT or single arginine methylation site mutants. Bars represent mean. At least 300 cells over three biological replicates of each strain were analyzed. *, *P = 0.0351; ****, P = <0.0001*. **(F)** α-strep western blot of L5::*sepIVA-*strep WT or single arginine methylation site mutants. mc^2^155 without a strep tag was used as a negative control (lane 2). The non-specific band serves as a loading control. P-values were calculated using ordinary one-way ANOVA, Dunnett’s multiple comparisons test, with a single pooled variance in GraphPad Prism (v9.2). At least two western blots were performed for each strain shown, and representative results are displayed.

Our analysis of clustered arginine mutants of *sepIVA* (Fig. 1AB) identified two sets of mutants with interesting phenotypes. If the lysine and methionine mutants mimic the different arginine methylation states, and if that methylation has a physiological role, then we expect to see opposite phenotypes in the methyl-ablative (R->K) and methyl-mimetic (R->M) strains compared to the wild-type. The *sepIVA* R3K, R19K, R31K mutant, hereafter called the NT-K mutant, as these are the most N-terminal arginines, exhibited slightly faster growth, and shorter cells than the wild-type, while the NT-M mutant (*sepIVA* R3M, R19M, R31M) had slower growth (Fig. 1AB). The mutant *sepIVA* R224K, R228K, R230K, R234K, R245K, hereafter called the CT-K mutant, had slow growth and (Fig. 1A) very short cell length (Fig. 1B). The C-terminal methionine mutant (*sepIVA* R224M, R228M, R230M, R234M, R245M) was not viable, likely because the protein was unstable (Fig. 1C).

To examine whether individual arginine residues were responsible for these growth and cell length defects, we generated *Msmeg* strains with single methylation site mutations in the N-terminal and C-terminal arginines. We found that single methylation site mutations near the N-terminus of *sepIVA* did not have the same effects on growth and cell length as the triple mutant (Fig. 1DE). Since more than one methylation site mutant near the N-terminus was needed for these phenotypes to be apparent, we used the NT-K and NT-M triple mutants to study the potential role of arginine methylation on SepIVA’s N-terminus (Fig. 2,3,4,6). The single methylation site mutations on the C-terminus of SepIVA had different phenotypes from the quintuple CT-K mutant (Fig. 1ABDE). Both the *sepIVA* R230K and R230M exhibited slow growth and long cells (Fig. 1DE): this indicates that these mutations likely do not represent different methylation states and are instead both loss-of-function mutations. Another methylation site, R234, only showed growth defects in the methionine mutant (Fig. 1DE). Since the phenotypes of the C-terminus single site mutants were distinct from the phenotype of the CT-K quintuple mutant, we believe the short cell length seen in the CT-K mutant was a result of suppressor mutations.

**Figure 2.**
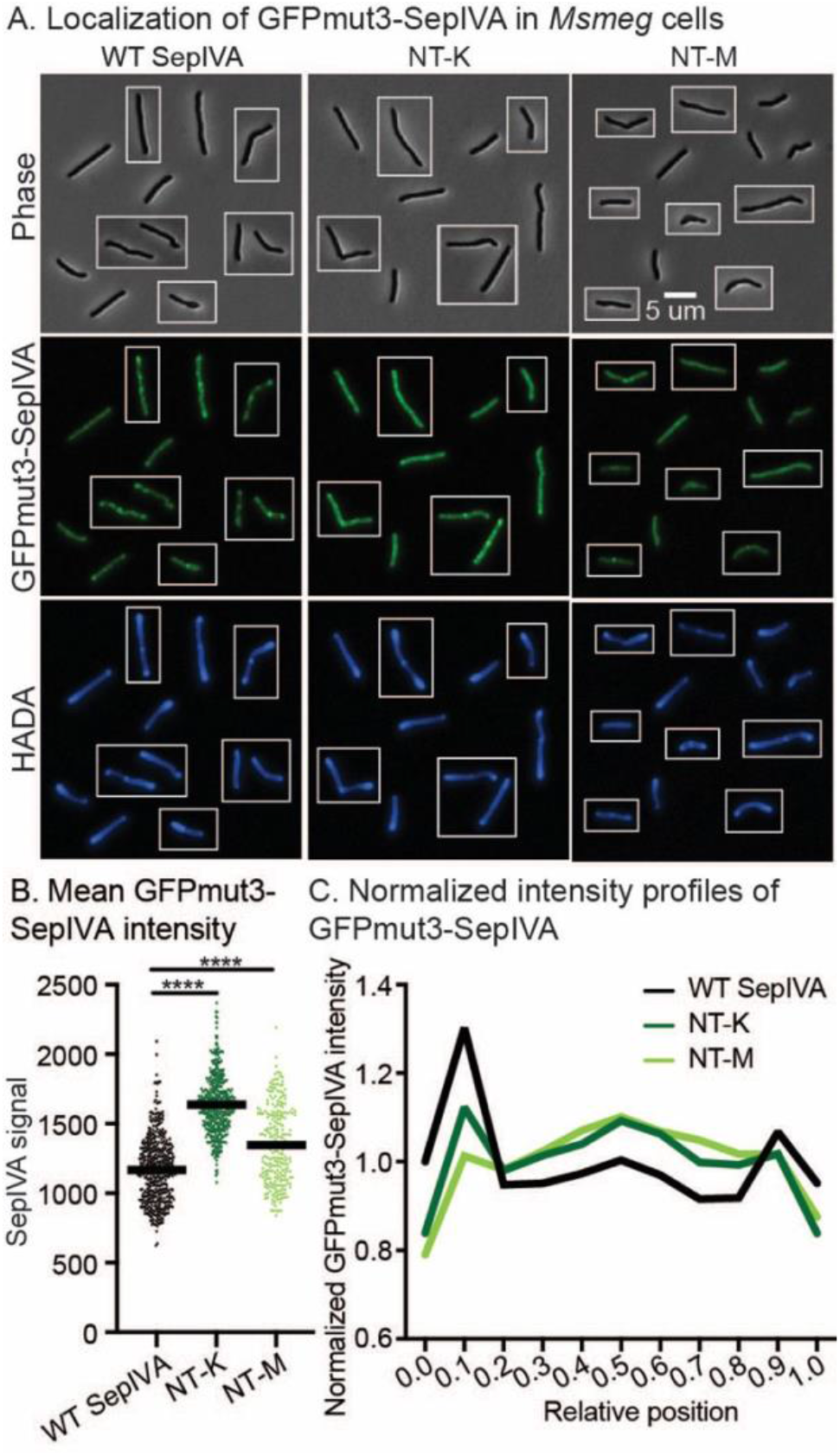
Methylation site mutations affect polar localization of SepIVA. **(A)** Images of *Msmeg* cells expressing *L5::GFPmut3-sepIVA* WT and methylation site mutants. Strains are merodiploid. Phase images are on the top row. Fluorescence images of GFPmut3-SepIVA are in the middle panel. HADA images are on the bottom row. Scale bar applies to all images. Pictures of several cells from images processed identically were pasted together. **(B)** Mean intensity values per cell of cells expressing *L5::GFPmut3-sepIVA* WT and methylation mutants. At least 250+ cells across three biological replicates of each mutant were analyzed in MicrobeJ. Bar represents mean intensity value between cells. ****, *P = <0.0001*. P-values were calculated using ordinary one-way ANOVA, Dunnett’s multiple comparisons test, with a single pooled variance in GraphPad Prism (v9.2). **(C)** Normalized Intensity profiles of GFP-SepIVA signal of WT and methylation site mutants. The solid line represents the mean intensity value across the relative position of the cell. Normalized intensity values were generated by dividing each position’s intensity value by the mean intensity value. Cells were pole-sorted such that the brighter pole in the HADA channel was set to 0 on the X axis.

**Figure 3.**
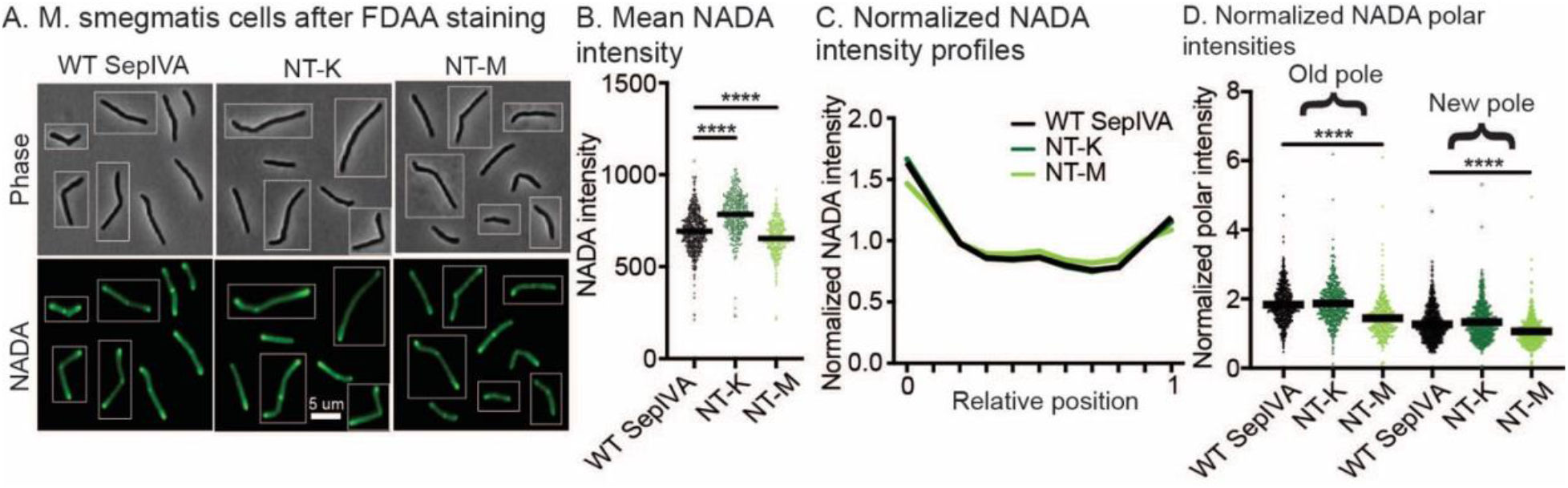
Methylation site mutations of SepIVA affect peptidoglycan metabolism. **(A)** Images of M. smeg cells expressing *L5::sepIVA* WT and methylation site mutants. Scale bar applies to all images. Pictures of several cells from images processed identically were pasted together. **(B)** Mean NADA intensity values of *M. smeg* cells expressing *L5::sepIVA* WT and methylation site mutants. At least 250+ cells across three biological replicates of each mutant were analyzed in MicrobeJ. Bar represents mean intensity value. ****, *P = <0.0001*. P-values were calculated using ordinary one-way ANOVA, Dunnett’s multiple comparisons test, with a single pooled variance in GraphPad Prism (v9.2). **(C)** Normalized intensity profiles of NADA signal of WT and methylation mutants. The solid line represents the mean intensity value across the relative position of the cell. Intensity values were normalized by dividing each strain’s intensity value by its mean intensity value. Cells were pole-sorted such that the brighter pole in the NADA channel was set to 0 on the X axis. **(D)** Normalized polar NADA intensities of WT and methylation site mutants. Intensity values were normalized by dividing each cell’s maximum polar intensity value by the mean intensity of the whole cell. ****, *P = <0.0001*. P-values were calculated using ordinary one-way ANOVA, Dunnett’s multiple comparisons test, with a single pooled variance in GraphPad Prism (v9.2).

**Figure 4.**
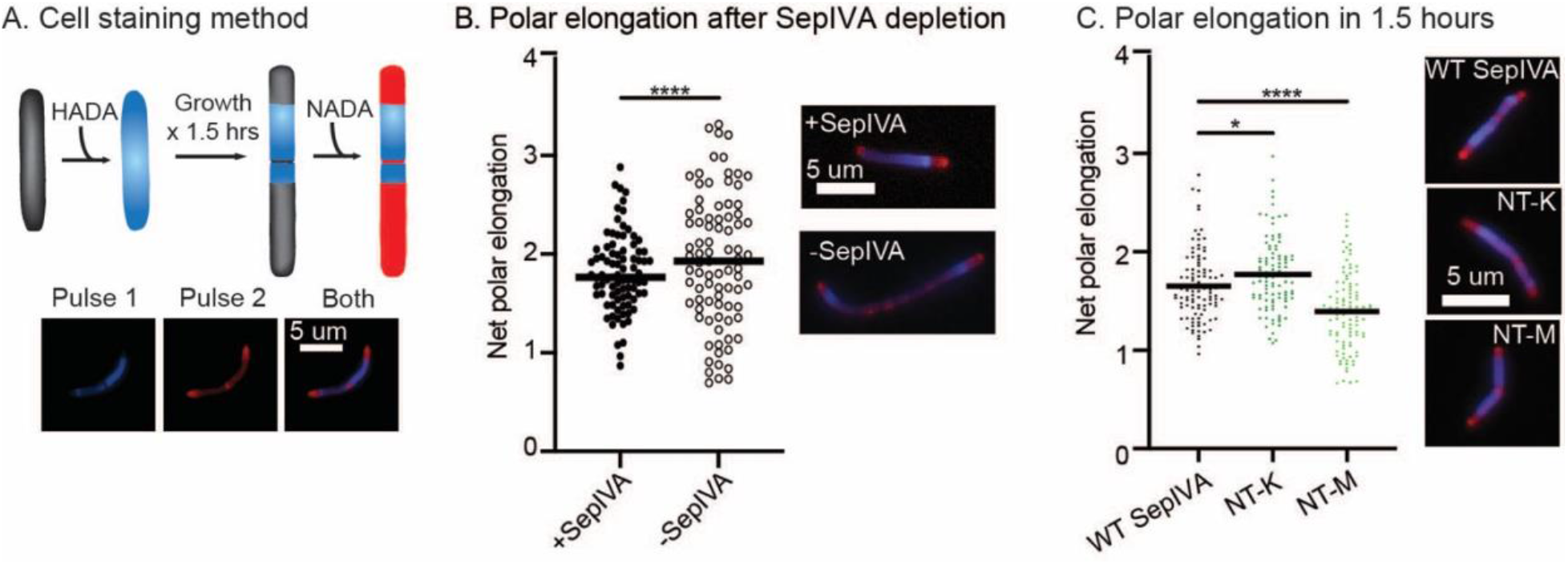
Methylation site mutations on SepIVA affect polar elongation. **(A)** Schematic of pulse-chase-pulse staining method. We stained growing cells with blue peptidoglycan dye HADA, then returned them to media for another 1.5 hours of growth, then stained them again with green dye NADA. We false-colored the green signal as red in these images to make the contrast clearer. **(B)** Length of poles after SepIVA depletion. +SepIVA (black dots) represent cells expressing SepIVA, -SepIVA (black-outlined dots) represent cells with depleted SepIVA protein. At least 100 cells across three biological replicates were analyzed. Bar represents median. Scale bar applies to all images. ****, *P = <0.0001*. P-value calculated by unpaired t-test using GraphPad Prism (v9.2). **(C)** Length of poles elongation after 1.5 hours in cells expressing *sepIVA* WT and methylation site allele swap mutants. At least 100 cells across three biological replicates were analyzed. Bar represents mean polar elongation. *, *P = 0.0461*, ****, *P = <0.0001*. P-values were calculated using ordinary one-way ANOVA, Dunnett’s multiple comparisons test, with a single pooled variance in GraphPad Prism (v9.2).

**Figure 5.**
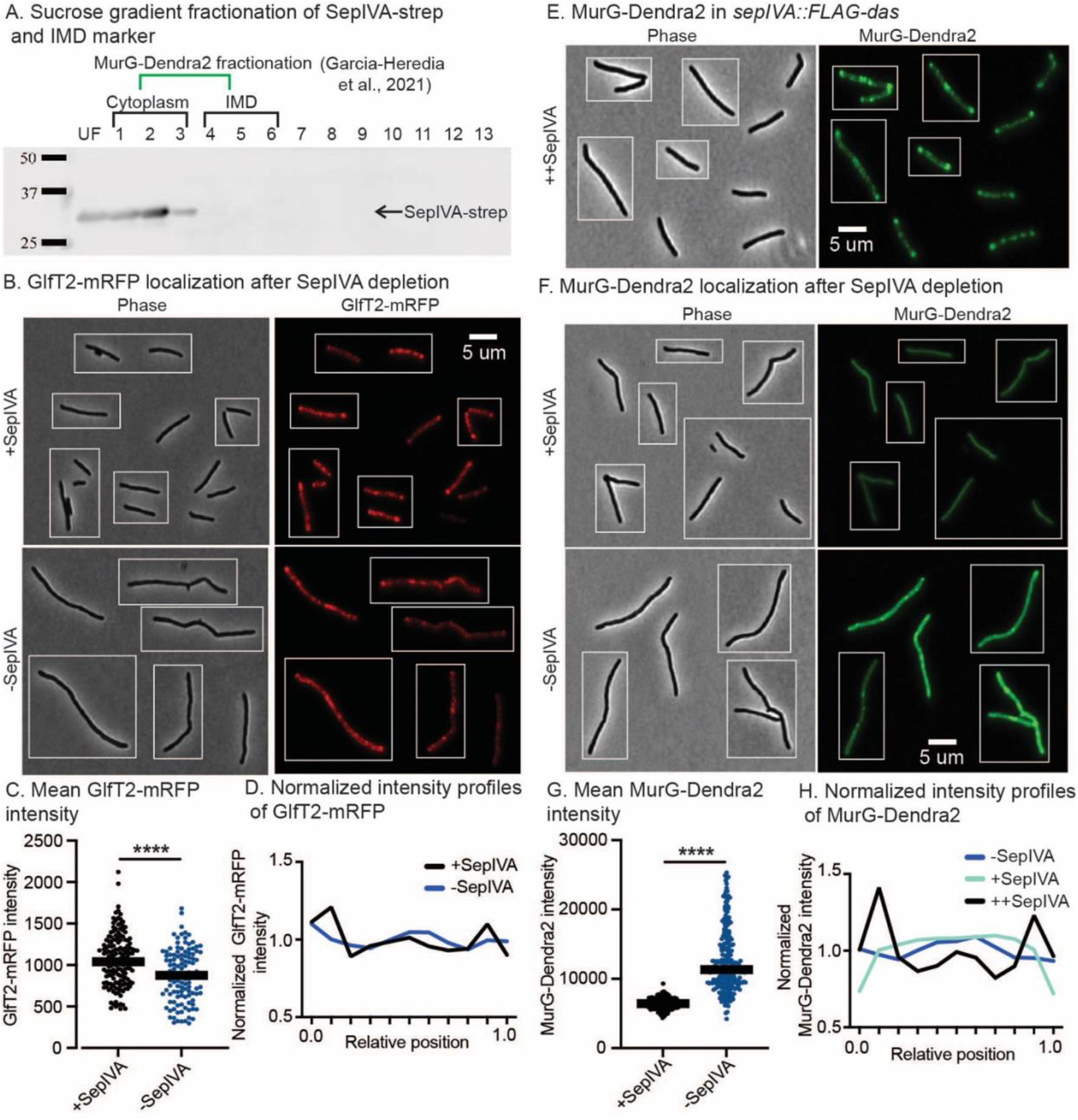
SepIVA has the same membrane association as MurG, and it affects MurG localization. **(A)** anti-Strep western blot of sucrose gradient fractions. Fractions 4-6 contain intracellular membrane domain (IMD) proteins. The green bracket denotes fractionation of MurG-Dendra2, as shown in Garcia-Heredia *et al*. 2021. **(B)** Micrographs of cells expressing or depleting SepIVA and induced for GlfT2-mRFP. Scale bar is 5 microns and applies to all images. Pictures of several cells from images processed identically were pasted together. **(C)** Mean GlfT2-mRFP intensity values of *Msmeg* cells expressing or depleting SepIVA and induced for GlfT2-mRFP. At least 250+ cells across three biological replicates of each mutant were analyzed in MicrobeJ. Bar represents mean intensity value. ****, *P = <0.0001*. P-value calculated by unpaired t-test using GraphPad Prism (v9.2). **(D)** Normalized intensity profiles of GlfT2-mRFP signal of cells expressing and depleting SepIVA and induced for GlfT2-mRFP. The solid line represents the mean intensity value across the relative position of the cell. Cells were pole-sorted such that the brighter pole in the HADA channel was set to 0 on the X axis. **(E)** Micrographs of cells expressing *sepIVA::FLAG-das*. This strain lacks inducible *sspB*, therefore, SepIVA cannot be depleted in this strain. We refer to this strain as ++SepIVA. These images are 12-bit. Scale bar is 5 microns and applies to all images. Pictures of several cells from images processed identically were pasted together. **(F)** Micrographs of cells expressing or depleting SepIVA while constitutively expressing MurG-Dendra2. These images are 16-bit. Scale bar is 5 microns and applies to all images. Pictures of several cells from images processed identically were pasted together. **(G)** Mean MurG-Dendra2 intensity values of *Msmeg* cells expressing or depleting SepIVA while constitutively expressing MurG-Dendra2. At least 250+ cells across three biological replicates of each mutant were analyzed in MicrobeJ. Bar represents mean intensity value. ****, *P = <0.0001*. P-value calculated by unpaired t-test using GraphPad Prism (v9.2). **(H)** Normalized intensity profiles of MurG-Dendra2 signal of cells expressing and depleting SepIVA. The solid line represents the mean intensity value across the relative position of the cell. Cells were pole-sorted such that the brighter pole in the HADA channel was set to 0 on the X axis.

**Figure 6.**
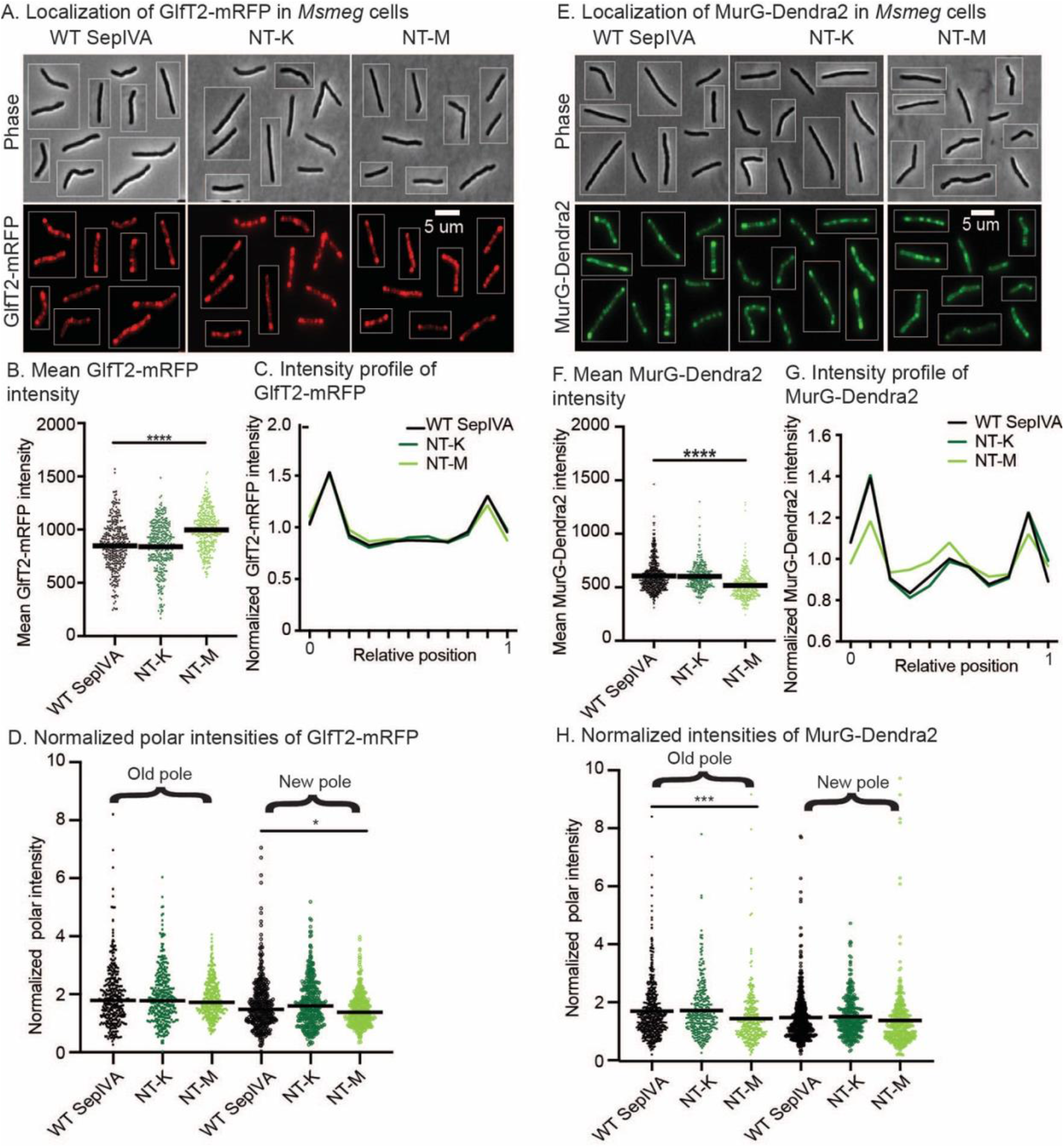
Methylation site mutations on SepIVA polar localization of MurG. **(A)** Images of *Msmeg* cells expressing *L5::sepIVA* WT or methylation site mutants and induced for GlfT2-mRFP. Scale bar is 5 microns and applies to all images. Pictures of several cells from images processed identically were pasted together. **(B)** Mean GlfT2-mRFP intensity values of *Msmeg* cells expressing *L5::sepIVA* WT or methylation site mutants and induced for GlfT2-mRFP. At least 250+ cells across three biological replicates of each mutant were analyzed in MicrobeJ. Bar represents mean intensity value. ****, *P = <0.0001*. **(C)** Normalized intensity profiles of GlfT2-mRFP signal of cells expressing *L5::sepIVA* WT or methylation site mutants and induced for GlfT2-mRFP. The solid line represents the mean intensity value across the relative position of the cell. Cells were pole-sorted such that the brighter pole in the HADA channel was set to 0 on the X axis. **(D)** Normalized polar GlfT2-mRFP intensities of WT and methylation site mutants. Intensity values were normalized by dividing each cell’s maximum polar intensity value by the mean intensity of the whole cell. *, *P = 0.0111*. **(E)** Images of *Msmeg* cells expressing L5::*sepIVA* WT or methylation site mutants and constitutively expressing MurG-Dendra2. Scale bar is 5 microns and applies to all images. Pictures of several cells from images processed identically were pasted together. **(F)** Mean MurG-Dendra2 intensity values of *Msmeg* cells expressing *L5::sepIVA* WT and methylation site mutants. At least 250+ cells across three biological replicates of each mutant were analyzed in MicrobeJ. Bar represents mean intensity value. ****, *P = <0.0001*. **(G)** Normalized intensity profiles of MurG-Dendra2 signal of cells expressing *L5::sepIVA* WT and methylation site mutants. The solid line represents the mean intensity value across the relative position of the cell. Cells were pole-sorted such that the brighter pole in the HADA channel was set to 0 on the X axis. **(H)** Normalized polar MurG-Dendra2 intensities of WT and methylation site mutants. Intensity values were normalized by dividing each cell’s maximum polar intensity value by the mean intensity of the whole cell. ***, *P = 0.0002*. P-values were calculated using ordinary one-way ANOVA, Dunnett’s multiple comparisons test, with a single pooled variance in GraphPad Prism (v9.2).

We made all the mutants described above with C-terminal strep tags so we could test for protein stability using western blotting. (Fig. 1CF). To test the stability of mutant proteins that did not support growth in allele swap strains (Fig. 1A – missing bars), we transformed the vectors carrying the mutant *sepIVA*-strep into wild-type *Msmeg* with the wild-type *sepIVA* allele at its native locus to support growth. Using these merodiploid strains, we found that SepIVA-strep alanine mutant proteins that did not support growth were unstable (Fig. 1C, ‘*’ indicates merodiploid strain). Several of the mutants had significantly higher levels of SepIVA than the wild-type, including the NT-K, NT-M and CT-K mutants (Fig. 1C) which exhibited phenotypes. To test whether high levels of these mutant proteins might be responsible for the slow growth, we tested how overexpression of the CT-K mutant affected growth rates (Fig. S3). We found that overexpression of SepIVA CT-K did not slow growth in both WT *sepIVA* and *sepIVA* CT-K genetic backgrounds. This result shows that the increased protein levels are not responsible for the slow growth and short cell length of the *sepIVA* CT-K mutant. Other variants of SepIVA-strep are stable; though protein levels between mutants vary (Fig. 1CF), these variations do not correlate with phenotypes (Fig. 1ABDE).

These data suggest that methylation sites on SepIVA appear to play a role in how SepIVA regulates growth and division. Near the N-terminus, methyl-mimetic site mutations cause slow growth and lead to slightly shorter cells (Fig. 1). However, near the C-terminus of SepIVA, both R230 mutants are defective in division. At R234 the methyl-mimetic mutation results in slow growing, long cells, while the methyl-ablative mutation results in cells like the WT. For the remainder of this work, we focus on the N-terminal triple mutants and the single site mutants of R234.

According to TnSeq experiments and our previous work, *sepIVA* is an essential gene (DeJesus *et al*., 2017; Wu *et al*., 2018; Dragset *et al*., 2019). In another recent study, a mutant of *sepIVA* missing only residues 156-245, the C-terminus, was characterized and the authors described cell division defects (Pickford *et al*., 2020). We generated this C-terminus truncation mutant using L5 allele swapping and were able to generate colonies on plates. However, once in liquid culture, these mutants exhibited highly variable or no growth (Fig. S4). We interpret this to mean that the *sepIVA* C-terminus is largely essential, but that suppressor mutants can form easily to allow growth in its absence.

### Methylation site mutations on SepIVA affect its polar localization

We sought to determine if alterations at the arginine methylation sites could affect SepIVA’s localization. SepIVA normally localizes at the poles and septum, with stronger localization to the old pole, and spotty localization along the lateral walls in a pattern (Wu *et al*., 2018) that is seen in proteins that associate with the Intracellular Membrane Domain (IMD) (Meniche *et al*., 2014; Hayashi *et al*., 2016). To test this, we made merodiploid strains of *Msmeg* with GFPmut3-SepIVA methyl-mutant alleles expressed from the L5 site. Strains carrying GFPmut3-*sepIVA* as the only allele of *sepIVA* were nonviable, so merodiploid strains were used. We grew these strains in log. phase, and stained the cells with the fluorescent D-amino acid HADA, which preferentially labels sites of new peptidoglycan synthesis, and allows us to identify the faster growing, old pole as the pole that stains more brightly (Kuru *et al*., 2012; Baranowski *et al*., 2018). Upon imaging the cells, we found that methylation site mutations on SepIVA affect its distribution and total intensity (Fig. 2). Specifically, both NT-K and NT-M mutants showed inhibited localization of SepIVA to the fast-growing pole (Fig. 2AC), and increased localization to the cell septum. This suggests that regulation of these methylation sites affects SepIVA distribution throughout the cell. Methylation site mutants also exhibit differences in GFPmut3-SepIVA intensity broadly (Fig. 2B), however, we cannot conclude whether mean GFPmut3-SepIVA signal is an indicator of protein levels within the cell. Our findings suggest that SepIVA’s localization to the cell poles is affected by changes in the methylation sites on the N-terminus of SepIVA. We cannot discount the possibility that GFPmut3-SepIVA localizes differently from the untagged protein, but these results show at least that these mutations can affect the distribution of the fusion in a merodiploid background.

### Methylation site mutations on SepIVA affect peptidoglycan metabolism

SepIVA’s essentiality for cell division and sequence similarity to DivIVA proteins suggests that it regulates the cell wall. To determine if the methylation sites on SepIVA impact metabolism of cell wall components, we stained the allele swap *sepIVA-strep* arginine-mutant strains with NADA (Kuru *et al*., 2015), which is incorporated into metabolically active peptidoglycan mostly by periplasmic amino acid exchange by L,D-transpeptidases (LDTs) (Kuru *et al*., 2012; Baranowski *et al*., 2018). We found that the N-terminal methyl-mimetic site mutant, NT-M, showed slightly decreased NADA signal at the cell poles compared to the wild-type (Fig. 3). We also found that the NT-K mutant showed increased, and the NT-M mutant showed decreased, overall NADA incorporation compared to the wild-type (Fig. 3B). To observe differences in distribution of NADA, we looked at NADA incorporation across the length of the cell. Based on normalized polar NADA intensities, there is less NADA incorporation at both poles in the NT-M mutant (Fig. 3CD). These data suggest SepIVA affects peptidoglycan metabolism, though they do not address whether these effects are direct or indirect. SepIVA had previously been shown to be essential for division (Wu *et al*., 2018), but these differences in polar NADA incorporation indicate that SepIVA may also be a regulator of polar peptidoglycan metabolism.

### Methylation site mutations on SepIVA affect polar elongation

To determine if the changes in NADA staining in the *sepIVA* methyl-mutants were due to alterations in LDT-mediated peptidoglycan remodeling (Baranowski *et al*., 2018) or due to insertion of new cell wall at the poles, we measured polar elongation in the *sepIVA* wild-type and mutant strains. We developed a method of measuring elongation using a pulse of HADA (blue), 1.5 hours of outgrowth in media, and another pulse of NADA (green) stain. We then measured the distance of the NADA-stained but HADA-clear poles, which represents the polar growth during the 1.5-hour outgrowth (Fig. 4A).

First, we used this method to determine if depletion of SepIVA affects cell elongation. SepIVA depletion leads to a slight increase in net polar elongation (Fig. 4B). Because cell division is inhibited in the SepIVA-depletion condition, the increase in net polar elongation could be due to the increased age of the poles in non-dividing cells: older poles in mycobacteria grow faster (Aldridge *et al*., 2012), so cells that are not creating new poles through division will have older poles overall. From these data we conclude that SepIVA is not essential for elongation.

To determine if methylation sites on SepIVA could affect elongation, we performed pulse-chase-pulse staining on *sepIVA-strep* wild-type and methylation site mutant allele swap strains. We observed increased elongation in the NT-K methyl-ablative mutants, while the NT-M methyl-mimetic mutant showed decreased polar elongation (Fig. 4C). Since our methyl-ablative and methyl-mimetic site mutations show opposing effects on polar elongation, this suggests that methylation of SepIVA at these sites regulates polar elongation. We also performed this elongation assay on single methylation site mutants (Fig. S5). We found that individual N-terminal arginines had a similar trend - with the lysine mutants exhibiting greater elongation than the methionine mutants – but none of the single mutants had a statistically significant effect. From these results, we conclude that methylation at the R3, R19 and R31 residues work together in regulating elongation. These results show that SepIVA is an elongation regulator; since it is not required for elongation (Fig. 4B) we suggest that its function may be to help toggle growth between the poles and septum.

### SepIVA affects MurG localization

The Intracellular Membrane Domain (IMD) is a membrane domain that can be separated by sucrose density centrifugation from the cytoplasmic fractions and from the PM-CW (plasma membrane attached to the cell wall) fraction, which contains PBPs and other periplasmic cell wall enzymes (Hayashi *et al*., 2016). Microscopy from our previous study showed that GFPmut3-SepIVA has a similar localization pattern as the IMD-associated arabinogalactan precursor enzyme GlfT2 (Wu *et al*., 2018). To determine if SepIVA is associated with the IMD, we performed sucrose gradient fractionation on an *Msmeg* strain with a C-terminal strep tag on SepIVA (Fig. 5A). We found that SepIVA was found in the cytoplasmic fractions 1-2, as well as fraction 3. GlfT2 was previously shown to be found restricted to the IMD in fractions 4-6 (Hayashi *et al*., 2018). MurG, the final cytoplasmic enzyme in PG precursor synthesis, which also has a typical IMD-localization pattern (Meniche *et al*., 2014; García-Heredia *et al*., 2021), is found partially in the IMD and partially in the cytoplasmic fractions (García-Heredia *et al*., 2021). Thus, SepIVA and MurG have some overlap in their density-gradient distribution. These results show that SepIVA is not associated strongly with the IMD; however, the localization pattern of GFPmut3-SepIVA indicates that it may associate indirectly or weakly.

Because other DivIVA homologs are involved in recruiting proteins to their site of activity (Marston and Errington, 1999; Halbedel and Lewis, 2019), we hypothesized that perhaps SepIVA could be involved in recruiting proteins to its locations near the IMD and at the septum. To test this, we looked at the effects of SepIVA depletion on localization of GlfT2-mRFP and MurG-Dendra2. We built an *Msmeg* strain where the only copy of *sepIVA* has a C-terminal DAS degradation tag, which targets a protein to be proteolyzed by ClpP when the SspB adaptor is expressed by adding an Atc inducer (Kim *et al*., 2011; Wu *et al*., 2018). We transformed this depletion strain with either of the localization constructs. We then added Atc to induce the degradation of SepIVA-DAS, or did not add Atc in the control, and imaged the cells. We found that depletion of SepIVA had mild effects on GlfT2-mRFP localization (Fig. 5B-D). However, we found that MurG-Dendra2 localization was highly sensitive to SepIVA levels. In the strain with SepIVA-DAS and no *sspB*, MurG-Dendra2 had a localization pattern similar to the normal IMD pattern that has been described before (Meniche *et al*., 2014; Hayashi *et al*., 2016; García-Heredia *et al*., 2021) (Fig. 5EH). When we transformed the *sspB* vector into this strain but did not induce *sspB* expression, we expect a moderate decrease in SepIVA levels due to leaky SspB expression, and we see complete delocalization of MurG-Dendra2 (Fig. 5GH). When we induce SspB to strongly deplete SepIVA-DAS, MurG-Dendra2 localizes in puncta that are randomly distributed around the cell (Fig. 5GH). We observed moderate changes in mean intensity of GlfT2-mRFP, and large changes in MurG-Dendra2 intensity between the SepIVA depleted and wild-type strains (Fig. 5CG); we do not know the significance of these changes, and here focus instead on the changes in distribution (Fig. 5H).

Because SepIVA has different effects on GlfT2-RFP and MurG-Dendra2, we conclude that SepIVA does not affect IMD structure globally and is not required for IMD-association of GlfT2. Our results indicate, however, that SepIVA could have a role modulating MurG’s localization and/ or association with the membrane.

### Methylation site mutants on SepIVA affect polar localization of MurG

We determined that depletion of SepIVA affects localization of MurG more strongly than GlfT2 (Fig. 5). To determine whether arginine methylation site mutations in *sepIVA* affect MurG or GlfT2 localization, we imaged each fluorescent protein fusion in *sepIVA* wild-type and methyl-mutant strains allele swap strains. We found that the distribution of GlfT2-mRFP around the cell was unaffected by methylation site mutations (Fig. 6ABCD). However, MurG-Dendra2 localization was less polar and more septal in the methyl-mimetic NT-M mutant (Fig. 6EFGH) compared to the methyl-ablative mutant and the WT. This suggests that an unmethylated N-terminus of SepIVA may stimulate MurG’s association with the growing pole (Fig 6GH), possibly by recruiting it to that site (Fig. 2) to promote polar PG synthesis and polar elongation (Fig. 3,4). SepIVA with a methylated N-terminus may instead decrease MurG’s association to the poles (Fig. 6GH), which results in decreased polar peptidoglycan metabolism (Fig. 3CD) and elongation (Fig. 4C) resulting in slower growth (Fig. 1A). We are observing a correlation between MurG polar association (Fig. 6GH) and polar growth (Fig. 4C) in these strains, which could be due to SepIVA directly or indirectly regulating MurG.

To test if arginine methylation site mutants of *sepIVA* affect the FtsZ ring, we expressed *ftsZ-mcherry2b* in methylation mutants. Our results show that arginine methylation of SepIVA does not affect FtsZ-mcherry2B localization (Fig. S6).

### Residues near the C-terminus of SepIVA function in cell division

Our preliminary work on individual arginine mutations at the C-terminus of *sepIVA* indicated that these residues affect cell division (Fig. 1). We further characterized the *sepIVA* R234 methylation site mutants. We observed the methyl-ablative methylation site mutant, R234K, is like the WT in growth rate and cell length, while the methyl-mimetic R234M has slower growth and longer cell length compared to the WT (Fig. 7ABC). The *sepIVA* R234M cells also elongate slightly more than the WT (Fig. 7D), though this is likely to be from the increased age of poles in cells with inhibited cell division, as in Fig. 4B. Thus, the *sepIVA* R234M strain displays inhibited cell division. The R234M strain has higher net peptidoglycan metabolism, as measured by NADA staining, than the WT, while the R234K has lower peptidoglycan metabolism than the WT (Fig. 7E). Because the two R234 mutants have opposing phenotypes, they may mimic the methylated and unmethylated states of SepIVA at R234. The data from these mutants suggests that arginine methylation at R234 inhibits division, while the unmethylated divides like the WT, suggesting that in the conditions measured, SepIVA is likely not methylated at this residue. The *sepIVA* R234 mutant strains both have increased MurG-Dendra2 fluorescent signal, but comparable distributions to the wild-type. Here, MurG signal does not correlate with difference in cell wall metabolism or growth (Fig. 7BCDGH), suggesting that regulation of cell division through the C-terminus of SepIVA does not involve regulating MurG localization. Our data show that the R234 methylation site is crucial for SepIVA’s role in cell division.

**Figure 7.**
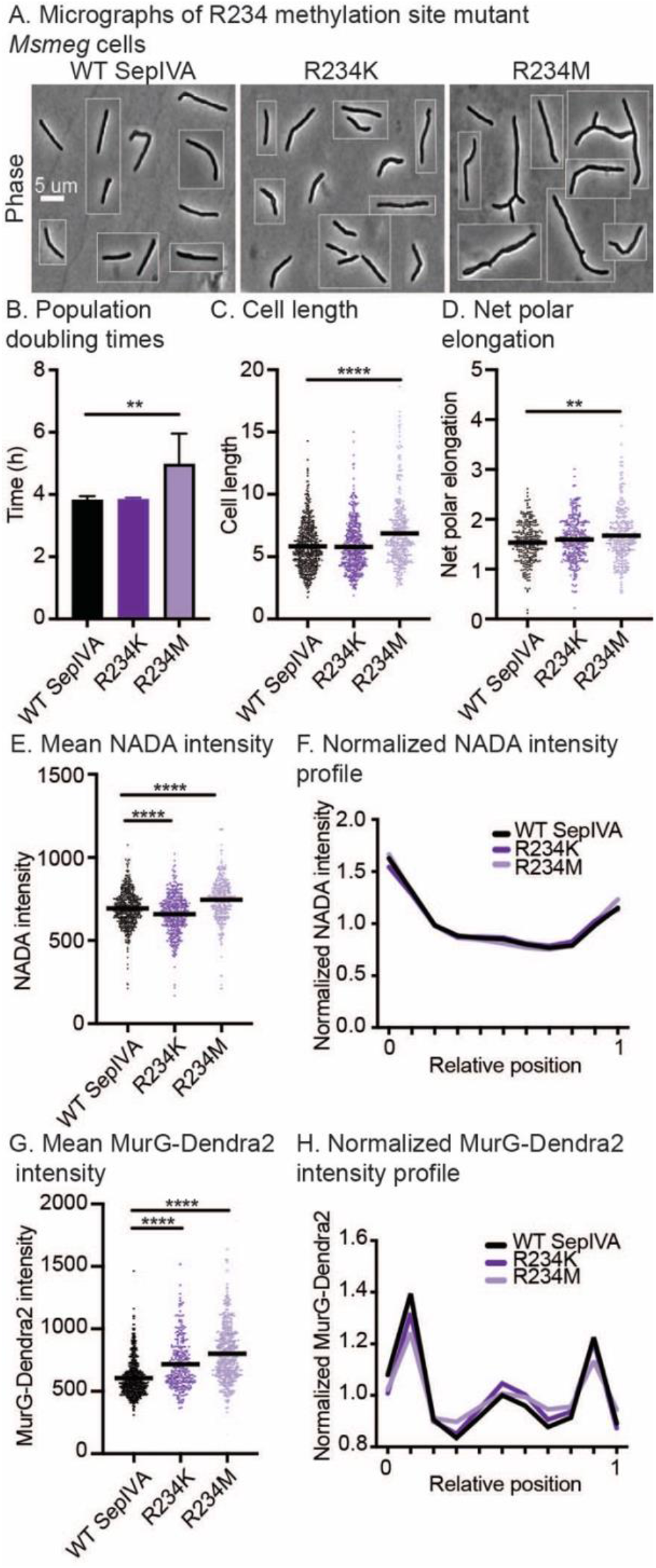
Methylation site R234 of SepIVA affects cell division. **(A)** Images of *Msmeg* cells expressing L5::*sepIVA* WT and R234 methylation site mutants. Scale bar is 5 microns and applies to all images. Pictures of several cells from images processed identically were pasted together. **(B)** Doubling time of *Msmeg* cells expressing WT or R234 methylation site mutants of SepIVA. No outline, darker color bars represent methyl-ablative methylation site mutants. Black outline bars represent methyl-mimetic methylation site mutants. At least 3 biological replicates of each strain were used. Error bars represent standard deviation. **, *P = 0.0012*. **(C)** Cell lengths of SepIVA WT or R234 methylation site mutants. Bars represent mean. At least 300 cells over three biological replicates of each strain were analyzed. ****, *P = <0.0001*. **(D)** Net polar elongation after 1.5 hours in cells expressing *L5::sepIVA* WT or R234 methylation site mutants. Bar represents mean polar elongation. At least 100 cells across three biological replicates were analyzed. **, *P = 0.0066*. **(E)** Mean NADA intensity values of *Msmeg* cells expressing *L5::sepIVA* WT or R234 methylation site mutants. Bar represents mean intensity value. At least 250+ cells across three biological replicates of each mutant were analyzed in MicrobeJ. *P = <0.0001*. **(F)** Normalized intensity profiles of NADA signal of cells expressing *L5::sepIVA* WT or R234 methylation site mutants. The solid line represents the mean intensity value across the relative position of the cell. Cells were pole-sorted such that the brighter pole in the NADA channel was set to 0 on the X axis. **(G)** Mean MurG-Dendra2 intensity values of *Msmeg* cells expressing *L5::sepIVA* WT or R234 methylation site mutants. Bar represents mean intensity value. At least 250+ cells across three biological replicates of each mutant were analyzed in MicrobeJ. ****, *P = <0.0001*. All P-values were calculated using ordinary one-way ANOVA, Dunnett’s multiple comparisons test, with a single pooled variance in GraphPad Prism (v9.2). **(H)** Normalized intensity profiles of MurG-Dendra2 signal of cells expressing *L5::sepIVA* WT or R234 methylation site mutants. The solid line represents the mean intensity value across the relative position of the cell. Cells were pole-sorted such that the brighter pole in the HADA channel was set to 0 on the X axis.

## Discussion

Although SepIVA is a DivIVA protein, the preliminary data on this factor indicate that its function is distinct from other described DivIVA proteins. All other described DivIVA proteins localize specifically to the curved membranes at poles and septa (Ramamurthi and Losick, 2009). SepIVA has some localization to the septum, but it mostly localizes in a ring around the cell near the pole, but not at the pole tip (Wu *et al*., 2018) (Fig. 2). This shows that SepIVA is not associated with primarily with curved membranes like other DivIVA proteins. Other described DivIVA proteins have been shown to have functions in promoting polar elongation (Kang *et al*., 2008; Letek *et al*., 2008; Hempel *et al*., 2008; Melzer *et al*., 2018), helping to inhibit cell division (Marston and Errington, 1999; Eswara *et al*., 2018) and/or to bind and regulate transcription factors (Eswaramoorthy *et al*., 2014), cell wall enzymes (Cleverley *et al*., 2019), cell division regulators (Eswara *et al*., 2018), and an envelope saccharide transporter (Hammond *et al*., 2022). None of these DivIVA proteins are essential for cell division, as SepIVA is (Wu *et al*., 2018). While some individual residues on DivIVA homologs have been characterized as being involved in these functions, none of these residues are conserved in SepIVA (Fig. S1). The low conservation between SepIVA and other DivIVA proteins, the difference in localization, and the wide variety of functions and binding partners across species make it impossible to use homology to predict the molecular functionality of SepIVA.

Mycobacteria must regulate peptidoglycan precursor synthesis in coordination with cell wall expansion and remodeling needs. Our data indicate that SepIVA likely helps regulate the peptidoglycan layer (Fig. 3) possibly partly through control of MurG (Fig. 5,6), which is the final enzyme in the construction of lipidII before it is flipped into the periplasm (Gee *et al*., 2012; Egan *et al*., 2020). In *E. coli*, MurG interacts with both elongation and division factors (Laddomada *et al*., 2016; Egan *et al*., 2020). According to our data, mild SepIVA depletion results in dispersion of MurG, and strong SepIVA depletion results in mislocalization of MurG (Fig. 5G-J). We speculate that SepIVA is somehow affecting MurG’s association with the Intracellular Membrane Domain (IMD) (Hayashi *et al*., 2016; García-Heredia *et al*., 2021) or other regulators that help cell wall metabolism transition between septal and elongative modes – our data do not indicate whether this regulation is direct or indirect. A third possibility is that SepIVA depletion is causing the type of cell wall damage that has previously been shown to cause MurG to re-localize to the lateral walls (Melzer *et al*., 2021). The arginine methylation sites on the N-terminus of SepIVA appear to toggle its effect on MurG localization in a way that correlates with polar elongation: the strain with the *sepIVA* N-terminal methyl-mimetic mutations have less polar growth (Fig. 4C) and less MurG at the poles (Fig. 6F). If SepIVA’s role in elongation regulation is to regulate MurG’s association and dispersal from the IMD domain at the poles - directly or indirectly - then it appears that when SepIVA is methylated at the N-terminus, it is more active in dispersing MurG from the poles.

Arginine methylation sites near the C-terminus of SepIVA, specifically R234, seem to play a role in regulating cell division (Fig. 7). The methyl-mimetic R234M mutant showed inhibited division (Fig. 7) while the methyl-ablative, R234K, did not. Methylation at this arginine residue could be inhibiting cell division. These effects on cell division revealed through mutations at the C-terminus appear to be independent of MurG localization (Fig. 7G).

Studies on protein arginine methylation done in eukaryotes support its importance in cell cycle progression (Biggar and Li, 2015; Raposo and Piller, 2018). Our data establish that protein arginine methylation sites on SepIVA affect the mycobacterial cell cycle as well. We have found that the arginine methylation sites on SepIVA affect both the elongation and division stages of the cell cycle. This work establishes protein arginine methylation as a physiologically important post-translational modification in bacteria and identifies SepIVA as a potential mediator between the elongation and division processes.

## Methods and Materials

### Bacterial strains and culture conditions

*Mycobacterium smegmatis* (mc^2^155) was cultured in 7H9 medium (Becton, Dickinson, Franklin Lakes, NJ) with 5 g/L bovine serum albumin (BSA), 2 g/L glucose, 0.85 g/L NaCl, 0.003 g/L catalase, 0.2% glycerol, and 0.05% Tween 80 or plated on LB agar. For *M. smeg* cultures, antibiotic concentrations were: 20 μg/mL nourseothricin, 20 μg/mL zeocin, 20 μg/mL streptomycin, 25 μg/mL kanamycin, 50 μg/mL hygromycin, and 30 μg/mL apramycin. For *E. coli* cultures, antibiotic concentrations were: 40 μg/mL nourseothricin, 25 μg/mL zeocin, 50 μg/mL streptomycin, 50 μg/mL kanamycin, 100 μg/mL hygromycin, and 50 μg/mL apramycin *E. coli* TOP10 or DH5α were used for cloning.

### Strain construction

Because it is essential in *M. smegmatis, sepIVA* mutants were constructed by first integrating an additional copy of *sepIVA* at the L5 integration site. After complementing *sepIVA* at the L5 site, *sepIVA* at its native locus was knocked out via recombineering (van Kessel and Hatfull, 2008), as previously described (Boutte *et al*., 2016). Mutant copies of *sepIVA* replaced wildtype *sepIVA* at the L5 site via allelic exchange (Pashley and Parish, 2003; Kieser *et al*., 2015).

*sepIVA* depletion strains were constructed as previously described (Wu *et al*., 2018). Merodiploid expression constructs of GlfT2-mRFP, FtsZ-mcherry2B, and MurG-Dendra2 were integrated at the Tweety integrase site (Pham *et al*., n.d.). The merodiploid construct expressing GFPmut3-SepIVA was integrated at the L5 integrase site. All strains used are listed in Table S3 of the supplementary material, along with plasmids listed in Table S4 and primers listed in Table S5.

### Growth rate assay

Biological replicates of *sepIVA* arginine mutants were grown to log phase in 7H9 medium. Cultures were diluted to OD_600_ = 0.1 in 200 μL 7H9 medium in a non-treated 96-well plate. Optical density at 600 nm was measured over the next 18-24 hours using a plate reader (BioTek Synergy neo2 multi-mode reader) at 37°C, shaking continuously. Population doubling times were determined using the exponential growth equation and least squares regression fitting method in GraphPad Prism (version 9.2). P values were calculated using ordinary one-way ANOVA, Dunnett’s multiple comparisons test, with a single pooled variance.

### Western blots

Log phase cultures (10 mL) were pelleted and resuspended in 500 μL PBS + 1 mM phenylmethylsulfonyl fluoride (PMSF). Cells were lysed using glass beads (MiniBeadBeater-16, model 607, Biospec). Supernatants from cell lysates were run on 4–12% NuPAGE Bis Tris precast gels (Life Technologies, Beverley, MA) using MES running buffer (Life Technologies, Beverly, MA). Proteins were transferred onto polyvinylidene difluoride (PVDF) membranes (GE Healthcare). Strep-tagged proteins were detected using rabbit anti-Strep-tag II antibody (1:1000, Abcam, ab76949) in Tris-buffered saline with Tween 20 (TBST) with 0.5% BSA and goat anti-rabbit IgG (H+L) horseradish peroxide (HRP)-conjugated secondary antibody (1:10,000, Thermo Fisher Scientific 31460) in TBST. Tagged proteins were visualized using chemiluminescent reagents (Thermo Fisher Scientific 34579).

### Microscopy

Cells were imaged using a Nikon Ti-2 widefield epifluorescence microscope with a Photometrics Prime 95B camera and a Plan Apo 100X, 1.45 -numerical-aperture objective. Green-fluorescent images for GFPmut3 or Dendra2 localization or NADA staining were taken with a 470/40nm excitation filter and a 525/50nm emission filter. Blue-fluorescent images for HADA staining were taken with a 350/50nm excitation filter and a 460/50nm emission filter. Red-fluorescent images for mRFP and mcherry2B localization were taken with a 560/40nm excitation filter and a 630/70 emission filter. Images were captured using NIS Elements software. Images were analyzed using FIJI and MicrobeJ (Ducret *et al*., 2016). For cell detection using MicrobeJ, parameters for width and area were set. V-snapping cells were cut at the septum so daughter cells could be detected as individual cells. Overlapping cells were excluded from analysis.

Cell lengths, mean intensities, maximum intensities, and minimum intensities of at least 250 cells per genotype were quantified using MicrobeJ. Mean intensity profiles were plotted using the “XStatProfile” plotting tool in MicrobeJ. P-values were calculated using ordinary one-way ANOVA, Dunnett’s multiple comparisons test, with a single pooled variance using GraphPad Prism (v9.2).

### Localization of cell wall proteins upon SepIVA depletion

For depletion of SepIVA, log phase cells (10 mL) were incubated (37°C) with 500 ng/mL Anhydrotetracycline (ATc) for 7-9 hours. For induction of GlfT2-mRFP, 1 ng/mL IVN were added, and cells were incubated (37°C) for 5 hours. Cells were fixed for microscopy. Cells were fixed for microscopy with 1.6% paraformaldehyde for 10 minutes and then washed and resuspended in PBS tween80. Microscopy images were analyzed in ImageJ and MicrobeJ. P-values calculated by unpaired t-test using GraphPad Prism (v9.2).

### Fluorescent staining

HADA (ThermoFisher) and NADA (ThermoFisher) fluorescent D-alanine amino acids were used to stain the peptidoglycan cell wall layer (Kuru *et al*., 2012). For PG analysis, NADA stain was added to 1 mL exponentially growing cells at a final concentration of 1 ug/mL for 3 minutes at room temperature before washing twice in PBST. For pulse-chase experiments, HADA (1 ug/mL) was added for 15 minutes before cells were washed twice in 7H9, incubated (37°C) for 1.5 hours, and stained again using NADA (1 ug/mL) for 3 minutes at room temperature. Cells were washed twice in PBST and imaged. Images were analyzed in ImageJ (NIH). The length of the poles and the length of the entire cell were measured manually for each cell.

### Sucrose Density Gradient Fractionation

Sucrose density gradient fractionation was performed as previously described (1{Citation}5). In brief, cells were grown to log phase and lysed by nitrogen cavitation. Lysate was then placed on top of a 20-50% sucrose gradient and centrifuged at 35,000 rpm for 6 hours at 4°C. Twelve one-mL fractions were collected and analyzed.

### Immunoprecipitation and mass spectrometry

We previously performed LCQ-MS/MS to identify peptides from *Mtb* cell lysates (Garces *et al*., 2010). Here, we re-analyzed those data to search for modifications in peptides, using Andromeda software.

SepIVA-strep was immunoprecipitated from CB1223. Cells were grown to late log. phase, pelleted and resuspended in PBS with PMSF at 1 mM, lysed by bead beating, and clarified by centrifugation. Immunoprecipitation was performed with MagStrep type 3 XT beads (Iba Life Sciences, Gottingen, Germany) according to the manufacturers protocol. Liquid was evaporated from the immunoprecipitants, and then digested by Trypsin/Lys-C Mix, Mass Spec Grade protease (Promega), according to the manufacturer’s protocols for reduction, alkylation and two-step in-solution digestion of proteolytically resistant proteins. The digested peptide solution was then cleaned by C18-Ziptips (Sigma Aldrich) according to the manufacturers protocol.

LC-MS analysis was performed on LTQ Velos pro mass spectrometer (Thermo Scientific, CA, USA) combined with a UHPLC (UltiMate 3000, Dionex, USA). The digested peptides were separated by a nano viper analytical C18 column. (Acclaim pepMap, 150 mm × 75μm, 2μm, 100 Å, Thermo Scientific, CA, USA). A 60 min gradient method was used to separate the digested peptides (0–3 min 4.0%B, 3–50 min 4.0– 50.0% B, 50–50.1 min 50–90% B,50.1–55 min 90% B, 55–55.1 min 90–4% B, 55.1–60 min 4% B; mobile phase A: 0.1% FA in water; mobile phase B: 0.1% FA in 95% acetonitrile, 5% water). 5μL of digested samples was injected at 300 nL/min flow. The nano-electrospray ionization (ESI) source used with a fixed spray voltage at 2.4 kV and a heated capillary temperature at 275 °C. A full MS spectrum obtained in normal scan mode from 350 to 2000 m/z mass range. The data-dependent acquisition was performed to get the MS/MS spectra for the 5 most abundant ions. The software Proteome Discoverer (Thermo Scientific, USA) was used to search and identify the peptides. LC-MS was performed at the Shimadzu Center for Advanced Analytical Chemistry at UT Arlington.

## Acknowledgements

This work was funded by grant R15GM131317 from the NIH. We thank Heather Lake who contributed to optimization of some experiments, and Arash Emami Saleh who aided in data processing.

