## Supplemental Figures 1-6, supplemental tables 3-5 for "Arginine methylation sites on SepIVA help balance elongation and septation of the cell wall in *Mycobacterium smegmatis*"

Cg= *Corynebacterium glutamicum*
Sc= *Streptomycin coelicolor*
Bsub= *B. subtillis*
Lm= *L. monocytogenes*

```
Msmeg_SepIVA      --------------------------------------MYRVFEALDELGA---------        13
Mtb_SepIVA        --------------------------------------VYRVFEALDELSA---------        13
Bsub_DivIVA       MPLTPNDIHNKTFTKS---FRGYDEDEVNEFLAQVRKDYEIVLRKKTELEAKVNELDERI        57
Sc_DivIVA         MPLTPEDVRNKQFTTVRL-REGYDEDEVDAFLDEVEAELTRLLRENEDLRAKLAAATRAA        59
Cg_Wag31          MPLTPADVHNVAFNKPPIGKRGYNEDEVDQFLDLVEDALVQFQEENEDLKQQVEELEAQV        60
Msmeg_Wag31       MPLTPADVHNVAFSKPPIGKRGYNEDEVDAFLDLVENELTRLIEENADLRQRVAELDQEL        60
Mtb_Wag31         MPLTPADVHNVAFSKPPIGKRGYNEDEVDAFLDLVENELTRLIEENSDLRQRINELDQEL        60
                                                           . .   :*

Msmeg_SepIVA      ------------------------------------------------------IVEEAR        19
Mtb_SepIVA        ------------------------------------------------------IVEEAR        19
Bsub_DivIVA       GHFANIEETL---NKSILVAQEAAEDVKRNSQK----EAKL-------------------        91
Sc_DivIVA         AQNQQNMRKPPEPPQDQQHQQGPPQHQQGPPQQGGMPQQGMRGPGAPVPAGISGPPQQQM        119
Cg_Wag31          AGGTSSAASSSTAGAATA---AASKSVDEAALRK-EIEEKLRSEYASKLDDASKAAQKAQ        116
Msmeg_Wag31       AAARSGAGASSQATSSIP-------------------------LYEPEPEPA--------        87
Mtb_Wag31         AA-GGGAGVTPQATQAIP-------------------------AYEPEPGKP--------        86

Msmeg_SepIVA      GVPMTAGCVVPRGDV-------------------------------LELIDDIKDAI---        45
Mtb_SepIVA        GVPMTAGCVVPRGDV-------------------------------LELIDDIKDAI---        45
Bsub_DivIVA       ------------------------------------------------------------        91
Sc_DivIVA         GGPMGGPPQLPSG----------APQLPAGPGGQGGPQGPGPMGQGPGPMGQGPGPMQGQ        169
Cg_Wag31          NDAKSAQDQLQRAQADAKAARDEAEKAKAEAKA--AASSNTTKAAAVGAVGAGTG-----        169
Msmeg_Wag31       ---------------------------PAPPQ-------PV-----------YEA-----        97
Mtb_Wag31         ------------------------------------------------------A-----        87

Msmeg_SepIVA      --PGE----LDDAQDVLDAR----DSLLREAKEHSESVISGANAEADSVLSHARAEADRL        95
Mtb_SepIVA        --PGE----LDDAQDVLDAR----DSMLQDAKTHADSMVSSATTEAESILNHARTEADRI        95
Bsub_DivIVA       -------------------------------------IVREAEKNADRIINESLSKSRKI        114
Sc_DivIVA         MGPGPMGGPMGGPQGPGGPGGPGMPGQGGPGGDSAARVLSLAQQTADQAIAEARSEANKI        229
Cg_Wag31          ---------------AAVATG----AANVDTHMQAAKVLGLAQEMADRLTSEARSESKSM        210
Msmeg_Wag31       ---------------PAQPAA----PQSEDTAVRAARVLSLAQDTADRLTSTAKAEADKL        138
Mtb_Wag31         ---------------PAAVSA----GMNEEQALKAARVLSLAQDTADRLTNTAKAESDKM        128
                                                       ::  *   *:     : :::  :

Msmeg_SepIVA      LADAKAQADRMVAEARQHSERMVTEAREEAARLAAAAKREYEASTGRAKAEADRLIENGN        155
Mtb_SepIVA        LSDAKAQADRMVSEARQHSERMVADAREEAIRIATAAKREYEASVSRAQAECDRLIENGN        155
Bsub_DivIVA       AME---------------------------------------------------------        117
Sc_DivIVA         VG----------------------------------------------------------        231
Cg_Wag31          LDEAREAAEKQIEEANSTSNRTLEDARANAEK-----------QIAEAQNRADTLVNEAD        259
Msmeg_Wag31       LSDARAQAEAMVSDARQTAE----------------------------------------        158
Mtb_Wag31         LADARANAEQILGEARHTAD----------------------------------------        148

Msmeg_SepIVA      ISYEKAIQEGIKEQQRLVSQTEIVATANAEATRLIDAAHAEADR----------------        199
Mtb_SepIVA        ISYEKAVQEGIKEQQRLVSQNEVVAAANAESTRLVDTAHAEADR----------------        199
Bsub_DivIVA       ------------------------------------------------------------        117
Sc_DivIVA         -------------------------EARSRAEGLERDARAKADALERDAQEKHRVAMGSL        266
Cg_Wag31          AKAKNLISEAEKKSAA------TLAASTSRAEAQIRQAEDKANALQADAERKHTETMAAV        313
Msmeg_Wag31       ----TTVSEARQRADA------MLADAQTRSEAQLRQAQEKADALQADAERKHSEIMGTI        208
Mtb_Wag31         ----ATVAEARQRADA------MLADAQSRSEAQLRQAQEKADALQADAERKHSEIMGTI        198

Msmeg_SepIVA      --------------------LRGECDIYVDSKLAEFE-----------------------        216
Mtb_SepIVA        --------------------LRGECDIYVDNKLAEFE-----------------------        216
Bsub_DivIVA       ----------IEELKKQSKVFRTRFQMLIEAQLDLLKNDDWDHLLEYEVDAVFEEK--E-        164
Sc_DivIVA         ESARATLERKVEDLRGFEREYRTRLKSYLESQLRQLETQADDSLAPPRTPATASLPPSPA        326
Cg_Wag31          KEQQNALETRIAELQTFEREYRTRLKSLLEGQLEELNARGSSA--PTN----------NK        361
Msmeg_Wag31       NQQRTVLEGRLEQLRTFEREYRTRLKTYLESQLEELGQRGSAA--PVD----------SS        256
Mtb_Wag31         NQQRAVLEGRLEQLRTFEREYRTRLKTYLESQLEELGQRGSAA--PVD----------SN        246
                                       * . .  :: :*  :

Msmeg_SepIVA      ---------EFLNGTLRSVGRGRHQLRTTAGTHDYVTR----------------------        245
Mtb_SepIVA        ---------EFLNGTLRSVGRGRHQLRTAAGTHDYAVR----------------------        245
Bsub_DivIVA       ------------------------------------------------------------        164
Sc_DivIVA         PSMAPAGASAPSYGGNQSMGGGPGQSGPSYGGQQQMSPAMTQPMAPVRPQGPSPMGQAPS        386
```

```
Cg_Wag31        PSGE--------------------------------------------------------        365
Msmeg_Wag31     ANSDASGFGQFNRGNN--------------------------------------------        272
Mtb_Wag31       --ADAGGFDQFNRGKN--------------------------------------------        260


Msmeg_SepIVA    ------------        245
Mtb_SepIVA      ------------        245
Bsub_DivIVA     ------------        164
Sc_DivIVA       PMRGFLIDEDDN        398
Cg_Wag31        ------------        365
Msmeg_Wag31     ------------        272
Mtb_Wag31       ------------        260
```

**Supplemental figure 1. Multiple sequence alignment of SepIVA from *Msmeg* and other DivIVA homologs.** Red- arginine residues on SepIVA and conserved in other homologs. Light Blue- F17 and R18 residues are crucial for membrane binding in *B. subtillis* (1). Green- R20 is crucial for DivIVA-RodA interaction in *C. glutamicum* (2). Orange- Phosphorylation by PknA/B on T73 in *Mtb* (3). '*' denotes a single, conserved residue. ':' denotes conservation of highly similar residues. '.' denotes conservation of weakly similar residues. Multiple sequence alignment was generated using CLUSTAL O(1.2.4)(6).

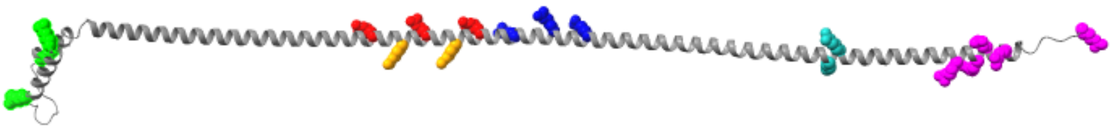

**Supplemental figure 2. Predicted SepIVA structure with colored arginine residues.** Tertiary structure of SepIVA protein as predicted by AlphaFold and visualized using UCSF ChimeraX. Arginine residues with methylations identified by mass spectrometry are grouped and colored to reflect fig. 1. We grouped arginines by their proximity to each other and their position on the SepIVA predicted structure. Arginine labels: green (R3, R19, R31; also referred to as the N-terminus mutants); red (R105, R116, R127); orange (R111, R122); dark blue (R134, R142, R149); teal (R199, R201); and pink (R224, R228, R230, R234, R245; also referred to as the C-terminus mutants).

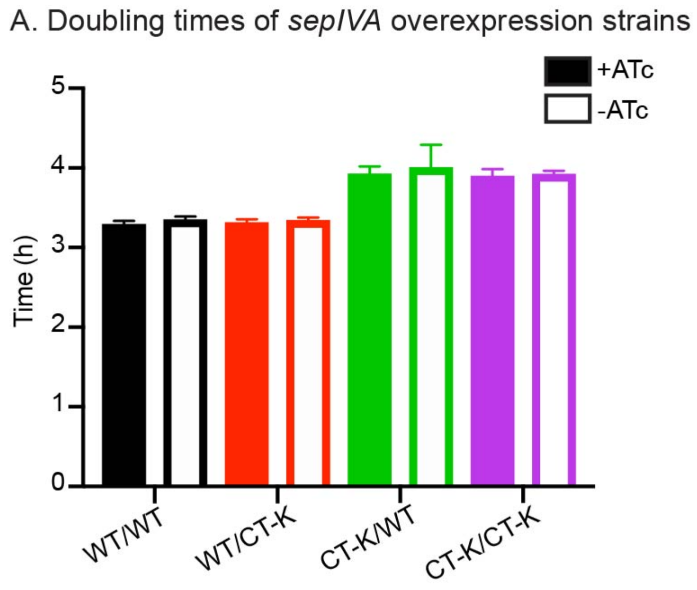


**Supplemental figure 3. Overexpression of *sepIVA* has no effect on population doubling time.** Overexpression of *sepIVA* was controlled by a tetracycline inducible promoter found on an episomal plasmid. +ATc denotes addition of anhydrotetracycline causing overexpression of SepIVA. Strains are labeled based on the *sepIVA* allele at the L5 integration site followed by the *sepIVA* allele overexpressed via episomal plasmid. At least two biological replicates were used per strain. Error bars represent standard deviation.

A. Gene arrangement of WT *sepIVA* and *sepIVA* C-terminus truncation mutant

B. Doubling times of WT *sepIVA* and *sepIVA* C-terminus truncation mutant

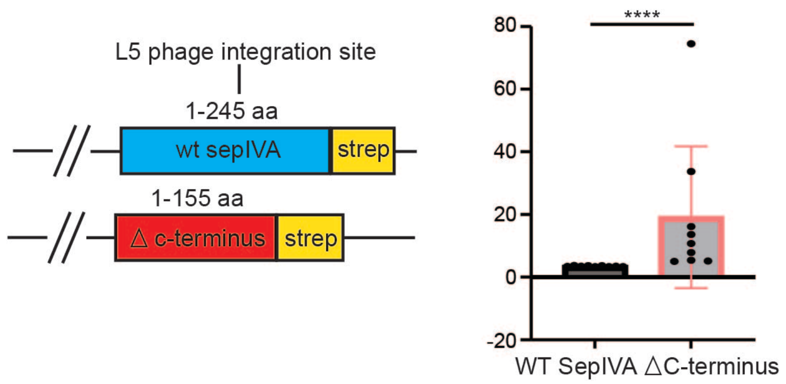


C. Growth chart of *sepIVA* C-terminus truncation mutants

| Isolate | Days before growth | Avg doubling time (h) |
|---|---|---|
| 1 | 7 | 39.971 |
| 2 | 4 | 16.13 |
| 3 | 8 | 6.492 |
| 4 | No growth | |
| 5 | 9 | 4.292 |
| 6 | No growth | |
| 7 | 3 | 33.7 |
| 8 | 11 | 44.2 |
| 9 | No growth | |
| 10 | No growth | |
| 11 | No growth | |
| 12 | 2 | 29.16 |
| 13 | 7 | 5.674 |
| 14 | No growth | |
| 15 | 7 | 13.6 |
| 16 | 5 | 5 |
| 17 | No growth | |
| 18 | 7 | 10.76 |
| 19 | No growth | |
| 20 | 7 | 19.75 |

**Supplemental figure 4. The C-terminus of SepIVA affects growth rate of *Msmeg*.** **(A)** Gene arrangement of WT *sepIVA* and *sepIVA* truncation mutant strains. The C-terminus truncation mutant only contains the first 155 amino acids of SepIVA. **(B)** Doubling times of WT *sepIVA* and *sepIVA* truncation mutants. Doubling times were only calculated for C-terminus isolates that were able to grow in liquid culture. At least 3 biological replicates of each strain were used. Error bars represent standard deviation. ****, *P = <0.0001.* P-value calculated by unpaired t-test using GraphPad Prism (v9.2). **(C)** Growth chart of *sepIVA* C-terminus truncation mutants. Twenty biological replicates were individually grown in liquid culture. “Days before growth” represents number of days rolling at 37C before visible growth occurred. Cultures that did not show visible growth after two weeks were deemed “no growth.” Doubling time was calculated only on isolates that grew in liquid culture.

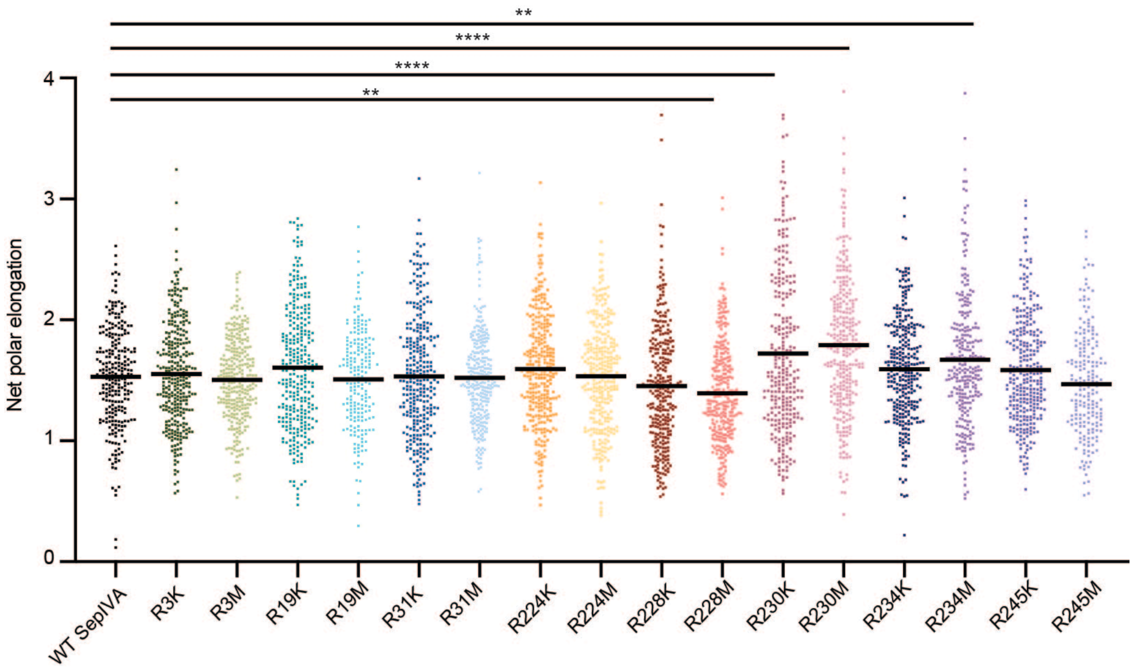


**Supplemental figure 5. Net polar elongation of single methylation site mutants.** Length of poles elongation after 1.5 hours in cells expressing *L5::sepIVA* WT and single methylation site mutants. At least 250 cells across three biological replicates were analyzed. Bar represents mean polar elongation. The adjusted p-values of the mutant compared to the WT are: R3M = 0.0009; R19M = 0.0066; and R31K = 0.0143*;* R31M = 0.0051; R224M = 0.0163; R228K = <0.0001; R228M = <0.0001; R230M = 0.0086; R234K = <0.0001; R245M = 0.0002. P-values were calculated using ordinary one-way ANOVA, Dunnett's multiple comparisons test, with a single pooled variance in GraphPad Prism (v9.2).

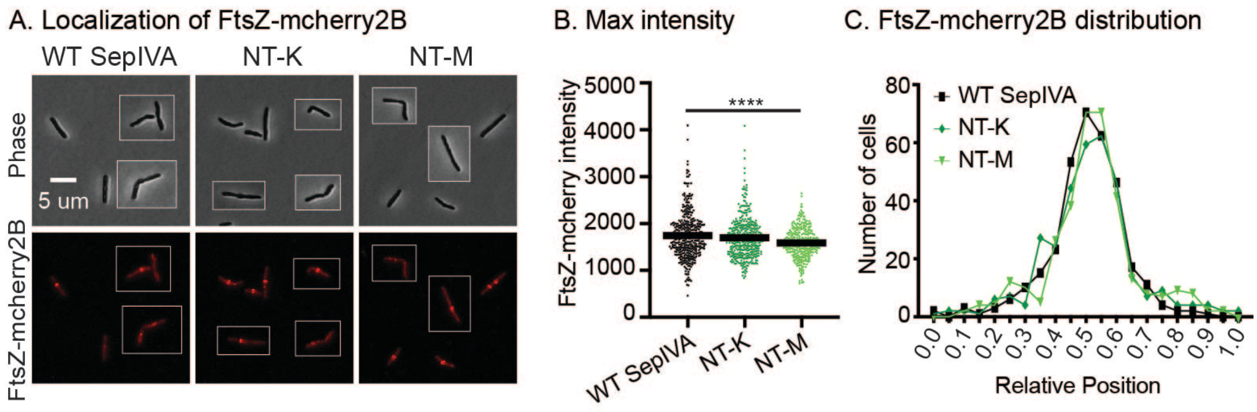


**Supplemental figure 6. Arginine methylation of SepIVA does not affect FtsZ-mcherry2B localization. (A)** Images of cells expressing L5::*sepIVA* WT and arginine methylation mutants and TW::*ftsZ-mcherry2B.* Pictures of several cells from images processed identically were pasted together. Scale bar applies to all images. **(B)** Max intensity values of 250+ cells expressing L5::*sepIVA* WT and arginine methylation mutants and TW::*ftsZ-mcherry2B.* Bar represents mean intensity value between cells. The adjusted p-values of each mutant compared to the WT are: NT-K = 0.2925; NT-M = <0.0001; and CT-K = 0.0436. At least 2 biological replicate strains were imaged for each genotype. P-values were calculated using ordinary one-way ANOVA, Dunnett's multiple comparisons test, with a single pooled variance in GraphPad Prism (v9.2). **(C)** Histogram of distribution FtsZ-mcherry2B max intensity value over the relative length of the cell. At least 250+ cells were analyzed. Relative positions of intensities were analyzed in MicrobeJ.

Supplemental Table 1. Re-analysis of mass spectrometry data of *Mtb* proteins to search for post-translational modifications.

Supplemental Table 2. Mass spectrometry data to search for post-translational modifications of SepIVA-strep from *Msmeg* lysates.

Supplemental Table 3. Strains.

| Strain # | Nickname | Genotype | Figure panel |
|---|---|---|---|
| CB1204 | | mc2155 ΔsepIVA::zeoR L5::pKK158-PsepIVA-sepIVA | |
| CB1223 | WT SepIVA | mc2155 ΔsepIVA::zeoR L5::pCB174- PsepIVA-sepIVA-strep | 1, 2, 3, 4C, 6, 7 |
| CB1486 | R134A R142A R149A | mc2155 ΔsepIVA::zeoR L5::pCB174-sepIVA-strep R134A R142A R149A | 1ABC |
| CB1510 | R134K R142K R149K | mc2155 ΔsepIVA::zeoR L5::pCB174-sepIVA-strep R134K R142K R149K | 1ABC |
| CB1502 | R111A R122A | mc2155 ΔsepIVA::zeoR L5::pCB174-sepIVA-strep R111A R122A | 1ABC |
| CB1489 | R111K R122K | mc2155 ΔsepIVA::zeoR L5::pCB174-sepIVA-strep R111K R122K | 1ABC |
| CB2141 | R105A R116A R127A* | mc2155 L5:: pCB174-sepIVA-strep R105A R116A R127A | 1C |
| CB1495 | R105K R116K R127K | mc2155 ΔsepIVA::zeoR L5::pCB174-sepIVA-strep R105K R116K R127K | 1ABC |
| CB1537 | NT-K or R3K R19K R31K | mc2155 ΔsepIVA::zeoR L5::pCB174-sepIVA-strep R3K R19K R31K | 1ABC, 2, 3, 4C, 6 |
| CB1561 | R3A R19A R31A* | mc2155 L5::pCB174-sepIVA-strep R3A R19A R31A | 1C |
| CB2126 | NT-M or R3M R19M R31M | mc2155 ΔsepIVA::zeoR L5::pCB174-sepIVA-strep R3M R19M R31M | 1ABC, 2, 3, 4C, 6 |

| | | | |
|---|---|---|---|
| CB1540 | R199A R201A | mc2155 ∆sepIVA::zeoR L5::pCB174-sepIVA-strep R199A R201A | 1 |
| CB1549 | R199K R201K | mc2155 ∆sepIVA::zeoR L5::pCB174-sepIVA-strep R199K R201K | 1 |
| CB1973 | CT-K or R224K R228K R230K R234K R245K | mc2155 ∆sepIVA::zeoR L5::pCB174-sepIVA-strep R224K R228K R230K R234K R245K | 1, 3, 4C |
| CB1565 | R224A R228A R230A R234A R245A* | mc2155 L5::pCB174-sepIVA-strep R224A R228A R230A R234A R245A | 1C |
| CB2143 | R224M R228M R230M R234M R245M* | mc2155 L5:: pCB174-sepIVA-strep R224M R228M R230M R234M R245M | 1C |
| CB2479 | R3K | mc2155 ∆sepIVA::zeoR L5::pCB174-sepIVA-strep R3K | 1DEF |
| CB2585 | R3M | mc2155 ∆sepIVA::zeoR L5::pCB174-sepIVA-strep R3M | 1DEF |
| CB1626 | R19K | mc2155 ∆sepIVA::zeoR L5::pCB174-sepIVA-strep R19K | 1DEF |
| CB2588 | R19M | mc2155 ∆sepIVA::zeoR L5::pCB174-sepIVA-strep R19M | 1DEF |
| CB2482 | R31K | mc2155 ∆sepIVA::zeoR L5::pCB174-sepIVA-strep R31K | 1DEF |
| CB2591 | R31M | mc2155 ∆sepIVA::zeoR L5::pCB174-sepIVA-strep R31M | 1DEF |
| CB2485 | R224K | mc2155 ∆sepIVA::zeoR L5::pCB174-sepIVA-strep R224K | 1DEF |
| CB2594 | R224M | mc2155 ∆sepIVA::zeoR L5::pCB174-sepIVA-strep R224M | 1DEF |
| CB2467 | R228K | mc2155 ∆sepIVA::zeoR L5::pCB174-sepIVA-strep R228K | 1DEF |
| CB2597 | R228M | mc2155 ∆sepIVA::zeoR L5::pCB174-sepIVA-strep R228M | 1DEF |
| CB2470 | R230K | mc2155 ∆sepIVA::zeoR L5::pCB174-sepIVA-strep R230K | 1DEF, 7 |
| CB2600 | R230M | mc2155 ∆sepIVA::zeoR L5::pCB174-sepIVA-strep R230M | 1DEF, 7 |
| CB2488 | R234K | mc2155 ∆sepIVA::zeoR L5::pCB174-sepIVA-strep R234K | 1DEF, 7 |
| CB2603 | R234M | mc2155 ∆sepIVA::zeoR L5::pCB174-sepIVA-strep R234M | 1DEF, 7 |
| CB2491 | R245K | mc2155 ∆sepIVA::zeoR L5::pCB174-sepIVA-strep R245K | 1DEF |
| CB2606 | R245M | mc2155 ∆sepIVA::zeoR L5::pCB174-sepIVA-strep R245M | 1DEF |
| CB2189 | GFP-WT SepIVA | mc2155 L5::pCB910-gfp-sepIVA | 2 |

| | | | |
|---|---|---|---|
| CB2190 | GFP-NT-K | mc2155 L5::pCB910-gfp-sepIVA R3K R19K R31K | 2 |
| CB2191 | GFP-NT-K | mc2155 L5::pCB910-gfp-sepIVA R3M R19M R31M | 2 |
| CB2137 | WT SepIVA/ ftsZ-mcherry | mc2155 ΔsepIVA::zeoR L5::pCB174-sepIVA-strep / pCB438-ftsZ-mcherry2B | S6 |
| CB2145 | NT-M/FtsZ-mcherry | mc2155 ΔsepIVA::zeoR L5::pCB174-sepIVA-strep R3M R19M R31M / pCB438-ftsZ-mcherry2B | S6 |
| CB2170 | NT-M/ftsZ-mcherry | mc2155 ΔsepIVA::zeoR L5::pCB174-sepIVA-strep R3K R19K R31K / pCB438-ftsZ-mcherry2B | S6 |
| CB2146 | CT-K/ftsZ-mcherry | mc2155 ΔsepIVA::zeoR L5::pCB174-sepIVA-strep R224K R228K R230K R234K R245K / pCB438-ftsZ-mcherry2B | S6 |
| CB2079 | OE WT/WT | mc2155 ΔsepIVA::zeoR L5::pCB174-sepIVA-strep /pCT56-pUV15-TetORH-sepIVA-strep | S3 |
| CB2082 | OE WT/CT-K | mc2155 ΔsepIVA::zeoR L5::pCB174-sepIVA-strep /pCT56-pUV15-TetORH-sepIVA-strep R224K R228K R230K R234K R245K | S3 |
| CB2085 | OE CT-K/WT | mc2155 ΔsepIVA::zeoR L5::pCB174-sepIVA-strep R224K R228K R230K R234K R245K/pCT56-pUV15-TetORH-sepIVA-strep | S3 |
| CB1965 | OE CT-K/CT-K | mc2155 ΔsepIVA::zeoR L5::pCB174-sepIVA-strep R224K R228K R230K R234K R245K/pCT56-pUV15-TetORH-sepIVA-strep R224K R228K R230K R234K R245K | S3 |
| CB1622 | SepIVA depletion, GlfT2 induction | mc2155 MSMEG_2416::FLAG-das-P38-orbit gi::pGMCgS-TetOFF-18-sspB TW::CT314-GlfT2-mRFP | 5BCDE |
| CB2257 | WT SepIVA/ GlfT2-mRFP | mc2155 ΔsepIVA::zeoR L5::pCB174-sepIVA-strep/pCT314-GlfT2-mRFP | 6ABC |
| CB2258 | NT-K/GlfT2-mRFP | mc2155 ΔsepIVA::zeoR L5::pCB174-sepIVA-strep R3K R19K R31K/ pCT314-GlfT2-mRFP | 6ABC |
| CB2259 | NT-M/GlfT2-mRFP | mc2155 ΔsepIVA::zeoR L5::pCB174-sepIVA-strep R3M R19M R31M/pCT314-GlfT2-mRFP | 6ABC |
| CB2287 | WT SepIVA/ MurG-Dendra2 | mc2155 ΔsepIVA::zeoR L5::pCT94-sepIVA-strep/pCT250-TwN-murG-dendra2 | 6DEF |
| CB2288 | NT-K/MurG-Dendra2 | mc2155 ΔsepIVA::zeoR L5::pCT94-sepIVA R3K R19K R31K-strep/ pCT250-TwN-murG-dendra2 | 6DEF |
| CB2289 | NT-M/MurG-Dendra2 | mc2155 ΔsepIVA::zeoR L5::pCT94-sepIVA R3M R19M R31M-strep/ pCT250-TwN-murG-dendra2 | 6DEF |
| CB2573 | R230K/MurG-Dendra2 | mc2155 ΔsepIVA::zeoR L5::pCT94-sepIVA R230K-strep/pCT250-TwN-murG-dendra2 | 7FH |
| CB2627 | R230M/MurG-Dendra2 | mc2155 ΔsepIVA::zeoR L5::pCT94-sepIVA R230M-strep/pCT250-TwN-murG-dendra2 | 7FH |
| CB2576 | R234K/MurG-Dendra2 | mc2155 ΔsepIVA::zeoR L5::pCT94-sepIVA R234K-strep/pCT250-TwN-murG-dendra2 | 7FH |
| CB2630 | R234M/MurG-Dendra2 | mc2155 ΔsepIVA::zeoR L5::pCT94-sepIVA R234M-strep/pCT250-TwN-murG-dendra2 | 7FH |
| CB2351 | SepIVA-DAS/ MurG-Dendra2 | mc2155 sepIVA::FLAG-das-P38-orbit/pCT250-TwN-murG-dendra2 | 5F |

| | | | |
|---|---|---|---|
| CB2352 | SepIVA depletion/MurG localization | mc2155 MSMEG_2416::FLAG-das-P38-orbit/pCT250-TwN-murG-dendra2 gi::pGMCgS-TetOFF-18-sspB | 5GHIJ |
| CB1166 | SepIVA depletion | mc2155 MSMEG_2416::FLAG-das-P38-orbit gi::pGMCgS-TetOFF-18-sspB | 4B |

Supplemental Table 4. Plasmid list.

| Strain # | Plasmid Name | Used in strains | Reference for parent vector |
|---|---|---|---|
| CB1199 | pKK158-PsepIVA-sepIVA | CB1204 | |
| CB1213 | pCB174-PsepIVA-sepIVA-strep | CB1223, CB1936, CB2079, CB2082, CB2137, CB2257 | (7) |
| CB1472 | pCB174-sepIVA-strep R134A R142A R149A | CB1486 | (7) |
| CB1513 | pCB174-sepIVA-strep R134K R142K R149K | CB1510 | (2) |
| CB1485 | pCB174-sepIVA-strep R111A R122A | CB1502 | (2) |
| CB1473 | pCB174-sepIVA-strep R111K R122K | CB1489 | (2) |
| CB1474 | pCB174-sepIVA-strep R105A R116A R127A | CB2141 | (2) |
| CB1475 | pCB174-sepIVA-strep R105K R116K R127K | CB1495 | (2) |
| CB1616 | pCB174-sepIVA-strep R3A R19A R31A | CB1561 | (2) |
| CB1617 | pCB174-sepIVA-strep R3K R19K R31K | CB1537, CB2170, CB2258 | (2) |
| CB2129 | pCB174-sepIVA R3M R19M R31M-strep | CB2126, CB2145, CB2259 | (2) |
| CB1618 | pCB174-sepIVA-strep R199A R201A | CB1540 | (2) |
| CB1619 | pCB174-sepIVA-strep R199K R201K | CB1549 | (2) |
| CB1620 | pCB174-sepIVA-strep R224A R228A R230A R234A R245A | CB1565 | (2) |
| CB1621 | pCB174-sepIVA-strep R224K R228K R230K R234K R245K | CB1965, CB2085 | (2) |

| | | | |
|---|---|---|---|
| CB2037 | pCB174-sepIVA-strep R224M R228M R230M R234M R245M | CB2143 | (2) |
| CB2462 | pCB174-PsepIVA-sepIVA-strep (R3K) | CB2479 | (2) |
| CB2550 | pCB174-PsepIVA-sepIVA-strep (R3M) | CB2585 | (2) |
| CB1666 | pCB174-PsepIVA-sepIVA-strep (R19K) | CB1647 | (2) |
| CB2551 | pCB174-PsepIVA-sepIVA-strep (R19M) | CB2588 | (2) |
| CB2463 | pCB174-PsepIVA-sepIVA-strep (R31K) | CB2483 | (2) |
| CB2552 | pCB174-PsepIVA-sepIVA-strep (R31M) | CB2591 | (2) |
| CB2464 | pCB174-PsepIVA-sepIVA-strep (R224K) | CB2485 | (2) |
| CB2553 | pCB174-PsepIVA-sepIVA-strep (R224M) | CB2594 | (2) |
| CB2442 | pCB174-PsepIVA-sepIVA-strep (R228K) | CB2467 | (2) |
| CB2554 | pCB174-PsepIVA-sepIVA-strep (R228M) | CB2597 | (2) |
| CB2443 | pCB174-PsepIVA-sepIVA-strep (R230K) | CB2470, CB2573 | (2) |
| CB2555 | pCB174-PsepIVA-sepIVA-strep (R230M) | CB2600, CB2627 | (2) |
| CB2465 | pCB174-PsepIVA-sepIVA-strep (R234K) | CB2488, CB2576 | (2) |
| CB2556 | pCB174-PsepIVA-sepIVA-strep (R234M) | CB2603, CB2630 | (2) |
| CB2466 | pCB174-PsepIVA-sepIVA-strep (R245K) | CB2491 | (2) |
| CB2557 | pCB174-PsepIVA-sepIVA-strep (R245M) | CB2606 | (2) |
| CB1694 | pCB910-gfp-sepIVA | CB2189 | |
| CB2193 | pCB910-gfp-sepIVA R3K R19K R31K | CB2190 | |
| CB2194 | pCB910-gfp-sepIVA R3M R19M R31M | CB2191 | |
| CB1957 | pCT56-pUV15-TetORH-sepIVA-strep | CB1936 | |
| CB1958 | pCT56-pUV15-TetORH-sepIVA-strep | CB1937 | |

| | | | |
|---|---|---|---|
| | R224K R228K R230K R234K R245K | | |
| CB1922 | pCT269-pMCtK-ftsZ-mcherry2B | CB1956, CB2137, CB2145, CB2146, CB2170 | |
| CB1036 | pGMCgS-TetOFF-18 sspB | CB1545, CB1622, CB1956 | (8) |
| CB2285 | pCT250-TwN-murG-dendra2 | CB2287, CB2288, CB2289, CB2351, CB2352, CB2573, CB 2576, CB2627, CB2630 | |
| CB942 | pCT258-sepIVA::FLAG-das-P38-orbit | CB1166, CB1622, CB2351 | (3) |

Supplemental Table 5. Primer list.

| Strain # | Feature | Primers |
|---|---|---|
| CB1213 | pL5::WT SepIVA | CTTTTTGCGTTTAATACTGCATGCACTCTAGAcgaggagttcgtcaagctct |
| | | CCTCTAGGGTCCCCAATTAATTAGCTAAAGCTTtcaCTTCTCGAACTGGGGGTG |
| CB1472 | pL5::SepIVA ARA | gaggccagcaccggcGCCgccaaggccgaggcc |
| | | aaggccgaggccgacGCGctcatcgagaacggc |
| | | gctggcctcgtactcGGCcttggctgccgccgc |
| CB1513 | pL5:SepIVA ARK | aaggccgaggccgacAAGctcatcgagaacggc |
| | | gctggcctcgtactcCTTcttggctgccgccgc |
| | | gaggccagcaccggcAAGgccaaggccgaggcc |
| CB1485 | pL5::SepIVA IRA | gtcgccgaggcgGCCcagcacagcgagcggatggtcaccgaggcgGCCgaggaagcggc |
| | | cggcgaaggagCCGgcggagccactggtaggcgagcgacacgacCCGgcggagccgctg |
| CB1473 | pL5::SepIVA IRK | gtcgccgaggcgAAGcagcacagcgagcggatggtcaccgaggcgAAGgaggaagcggc |
| | | cggcgaaggagGAAgcggagccactggtaggcgagcgacacgacGAAgcggagccgctg |
| CB1474 | pL5::SepIVA ORA | cagcacagcgagGCCatggtcaccgaggcgcgcgaggaagcggccGCCctcgcggcggc |
| | | ctgacgcgcctcggcgaccatGGCgtctgcctgggcttttgcatcgg |
| CB1475 | pL5::SepIVA ORK | cagcacagcgagAAGatggtcaccgaggcgcgcgaggaagcggccAAGctcgcggcggc |
| | | ctgacgcgcctcggcgaccatTTTgtctgcctgggcttttgcatcgg |
| CB1616 | pL5::SepIVA BFA | tcgaagaagcgGCCggtgtgccgatgaccgccggctgcgtcgtccctGCCggggatgtg |
| | | ccGGCcgcttcttcgacgatcgcgccgagctcgtcgagcgcttcaaaaacGGCgtacac |

| | | |
|---|---|---|
| CB1617 | pL5::SepIVA NT-K | gtgtacAAGgttttgaagcgctcgacgagctcggcgcgatcgtcgaagaagcgAAGgg |
| | | ccCTTcgcttcttcgacgatcgcgccgagctcgtcgagcgcttcaaaacCTTgtacac |
| | | tcgaagaagcgAAGggtgtgccgatgaccgccggctgcgtcgtccctAAGggggatgtg |
| CB1618 | pL5::SepIVA KA | cgacgcggcgcacgccgaggccgacGCCcttGCCggcgagtgcgacatctacgtcgaca |
| | | tgtcgacgtagatgtcgcactcgccGGCaagGGCgtcggcctcggcgtgcgccgcgtcg |
| CB1619 | pL5::SepIVA KK | cgacgcggcgcacgccgaggccgacAAActtAAGggcgagtgcgacatctacgtcgaca |
| | | tgtcgacgtagatgtcgcactcgccCTTaagTTTgtcggcctcggcgtgcgccgcgtcg |
| CB1620 | pL5::SepIVA LFA | ttctcaacgggaccctgGCCtcggtcggcGCCggaGCCcaccagttgGCCaccaccgcg |
| | | tcggcGCCggaGCCcaccagttgGCCaccaccgcggggacacacgactacgtgaccGCC |
| | | GGCggtcacgtagtcgtgtgtccccgcggtggtGGCcaactggtgGGCtccGGCgccga |
| | | GCCcaccagttgGCCaccaccgcggggacacacgactacgtgaccGCCCACTCCACCGGCGCCGA |
| | | TCGGCGCCGGTGGAGTGGGCggtcacgtagtcgtgtgtccccgcggtggtGGCcaactggtgGGC |
| CB1621 | pL5::SepIVA CT-K | cgcggtggtCTTcaactggtgCTTtccCTTgccgaccgaCTTcagggtcccgttgagaa |
| | | tcggcAAGggaAAGcaccagttgAAGaccaccgcggggacacacgactacgtgaccAAG |
| | | CTTggtcacgtagtcgtgtgtccccgcggtggtCTTcaactggtgCTTtccCTTgccga |
| | | AAGcaccagttgAAGaccaccgcggggacacacgactacgtgaccAAGCACTCCACCGGCGCCGA |
| | | TCGGCGCCGGTGGAGTGCTTggtcacgtagtcgtgtgtccccgcggtggtCTTcaactggtgCTT |
| CB2129 | pL5::SepIVA NT-M | acgcagccggcggtcatcggcacaccCATcgcttcttcgacgatcgcgccgagctcgtc |
| | | atgtcgtcgatgagttcgagcacatccccCATagggacgacgcagccggcggtcatcgg |
| | | ccgatgaccgccggctgcgtcgtccctATGgggatgtgctcgaactcatcgacgacat |
| | | tcgacgatcgcgccgagctcgtcgagcgcttcaaaacCATgtacacggcaacatcctc |
| CB2037 | pL5::SepIVA LFM | cacgtagtcgtgtgtccccgcggtggtCATcaactggtgCATtccCATgccgaccgaCATcagggtcccgttgagaa |
| | | tcacttCTCGAACTGGGGGTGGCTCCAGTCGGCGCCGGTGGAGTGCATggtcacgtagtcgtgtgtccccgcg |
| CB2462 | pL5::SepIVA R3K | aacaactaagcctggaggatgttgccgtgtacAAGgttttgaagcgctcgacgagctc |
| | | gagctcgtcgagcgcttcaaaacCTTgtacacggcaacatcctccaggcttagttgtt |
| CB2550 | pL5::SepIVA R3M | ctaagcctggaggatgttgccgtgtacATGgttttgaagcgctcgacgagctcggcgc |
| | | gcgccgagctcgtcgagcgcttcaaaacCATgtacacggcaacatcctccaggcttag |
| CB1666 | pL5::SepIVA R19K | ctcggcgcgatcgtcgaagaagcgAAGggtgtgccgatgaccgccggctgc |
| | | gcagccggcggtcatcggcacaccCTTcgcttcttcgacgatcgcgccgag |

| | | |
|---|---|---|
| CB2551 | pL5::SepIVA R19M | ctcggcgcgatcgtcgaagaagcgATGggtgtgccgatgaccgccggctgcgtcgtccc |
| | | gggacgacgcagccggcggtcatcggcacaccCATcgcttcttcgacgatcgcgccgag |
| CB2463 | pL5::SepIVA R31K | gtgccgatgaccgccggctgcgtcgtccctAAGggggatgtgctcgaactcatcgacga |
| | | tcgtcgatgagttcgagcacatccccCTTagggacgacgcagccggcggtcatcggcac |
| CB2552 | pL5::SepIVA R31M | ccgatgaccgccggctgcgtcgtccctATGggggatgtgctcgaactcatcgacgacat |
| | | atgtcgtcgatgagttcgagcacatccccCATagggacgacgcagccggcggtcatcgg |
| CB2464 | pL5::SepIVA R224K | tcgaggagtttctcaacgggaccctgAAGtcggtcggccgcggacgtcaccagttgcgg |
| | | ccgcaactggtgacgtccgcggccgaccgaCTTcagggtcccgttgagaaactcctcga |
| CB2553 | pL5::SepIVA R224M | gagttcgaggagtttctcaacgggaccctgATGtcggtcggccgcggacgtcaccagtt |
| | | aactggtgacgtccgcggccgaccgaCATcagggtcccgttgagaaactcctcgaactc |
| CB2442 | pL5::SepIVA R228K | tttctcaacgggaccctgcggtcggtcggcAAGggacgtcaccagttgcggaccaccgc |
| | | gcggtggtccgcaactggtgacgtccCTTgccgaccgaccgcagggtcccgttgagaaa |
| CB2554 | pL5::SepIVA R228M | tttctcaacgggaccctgcggtcggtcggcATGggacgtcaccagttgcggaccaccgc |
| | | gcggtggtccgcaactggtgacgtccCATgccgaccgaccgcagggtcccgttgagaaa |
| CB2443 | pL5::SepIVA R230K | aacgggaccctgcggtcggtcggccgcggaAAGcaccagttgcggaccaccgcggggac |
| | | gtccccgcggtggtccgcaactggtgCTTtccgcggccgaccgaccgcagggtcccgtt |
| CB2555 | pL5::SepIVA R230M | aacgggaccctgcggtcggtcggccgcggaATGcaccagttgcggaccaccgcggggac |
| | | gtccccgcggtggtccgcaactggtgCATtccgcggccgaccgaccgcagggtcccgtt |
| CB2465 | pL5::SepIVA R234K | tcggtcggccgcggacgtcaccagttgAAGaccaccgcggggacacacgactacgtgac |
| | | gtcacgtagtcgtgtgtccccgcggtggtCTTcaactggtgacgtccgcggccgaccga |
| CB2556 | pL5::SepIVA R234M | cggtcggtcggccgcggacgtcaccagttgATGaccaccgcggggacacacgactacgt |
| | | acgtagtcgtgtgtccccgcggtggtCATcaactggtgacgtccgcggccgaccgaccg |
| CB2466 | pL5::SepIVA R245K | accaccgcggggacacacgactacgtgaccAAGCACTCCACCGGCGCCGACTGGAGCCA |
| | | TGGCTCCAGTCGGCGCCGGTGGAGTGCTTggtcacgtagtcgtgtgtccccgcggtggt |
| CB2557 | pL5::SepIVA R245M | accaccgcggggacacacgactacgtgaccATGCACTCCACCGGCGCCGACTGGAGCCA |
| | | TGGCTCCAGTCGGCGCCGGTGGAGTGCATggtcacgtagtcgtgtgtccccgcggtggt |
| CB1694 | GFP::WT | ACACATGGCATGGATGAACTATACAAAGGTACCgtgtaccgagtttttgaagcgctcga |
| | | GGGTCCCCAATTAATTAGCTAAAGCTTctagcgggtcacgtagtcgtgtgtccccgcg |
| CB2193 | GFP::NT-K | ACATGGCATGGATGAACTATACAAAGGTACCgtgtacAAGgtttttgaagcgctcgacg |

| | | |
|---|---|---|
| | | GGGTCCCCAATTAATTAGCTAAAGCTTctagcgggtcacgtagtcgtgtgtccccgcg |
| CB2194 | GFP::NT-M | ACATGGCATGGATGAACTATACAAAGGTACCgtgtacATGgtttttgaagcgctcgacg |
| | | GGGTCCCCAATTAATTAGCTAAAGCTTctagcgggtcacgtagtcgtgtgtccccgcg |
| CB1936, CB1937 | WT OE, CT-K OE | GCCCGAAATGAGCACGATCCGCATGCTTAATatggtgtaccgagtttttgaagcgctcg |
| | | GTCCCCAATTAATTAGCTAAAGCTTGATtcacttCTCGAACTGGGGGTGGCTCCAGTCG |
| CB1922 | FtsZ-mcherry2B | gccctggaggagctcgggctgccggtgccgGGTTTCTTGTCAGTACGCGAAGAACCACG |
| | | ctcgacctgcaggcatgcaagcttGTCAGCGTAATGCTCTGCCAGT |
| CB1622 | GlfT2-mRFP | GGGTCCCCAATTAATTAGCTAAAGCTTTCACTTCTCGAACTGGGGGTGGCTCCAGTCGGC |
| | | AACGGAGGTACGCATATGGATATCGAATTCgaaagcagcaaagcatgagtgacatcccttccggcg |
| | | ctccttgatgaCGTCCTCGGAGGAGGCCATgagctctcgtccgactttctccggtgtctcggtgag |
| | | gagacaccggagaaagtcggacgagagctcATGGCCTCCTCCGAGGACGtcatcaaggag |
| CB2285 | MurG-Dendra2 | TCTAGGGTCCCCAATTAATTAGCTAAAGCTTCACTTGTCGTCGTCGTCCTTGTAGTCCT |
| | | ATGCTTAATTAAGAAGGAGATATACATatgatggtgaacaagggttcgcgggacaacaa |


1. Oliva MA, Halbedel S, Freund SM, Dutow P, Leonard TA, Veprintsev DB, Hamoen LW, Löwe J. 2010. Features critical for membrane binding revealed by DivIVA crystal structure. The EMBO Journal 29:1988–2001.

2. Sieger B, Bramkamp M. 2015. Interaction sites of DivIVA and RodA from Corynebacterium glutamicum. Front Microbiol 5.

3. Kang C-M. 2005. The Mycobacterium tuberculosis serine/threonine kinases PknA and PknB: substrate identification and regulation of cell shape. Genes & Development 19:1692–1704.

4. Rismondo J, Cleverley RM, Lane HV, Großhennig S, Steglich A, Möller L, Mannala GK, Hain T, Lewis RJ, Halbedel S. 2016. Structure of the bacterial cell division

determinant GpsB and its interaction with penicillin-binding proteins: *Listeria monocytogenes GpsB*. Molecular Microbiology 99:978–998.

5. Cleverley RM, Rutter ZJ, Rismondo J, Corona F, Tsui H-CT, Alatawi FA, Daniel RA, Halbedel S, Massidda O, Winkler ME, Lewis RJ. 2019. The cell cycle regulator GpsB functions as cytosolic adaptor for multiple cell wall enzymes. Nat Commun 10:261.

6. Fast, scalable generation of high-quality protein multiple sequence alignments using Clustal Omega | Molecular Systems Biology. https://www.embopress.org/doi/full/10.1038/msb.2011.75. Retrieved 27 January 2022.

7. Kieser KJ, Boutte CC, Kester JC, Baer CE, Barczak AK, Meniche X, Chao MC, Rego EH, Sassetti CM, Fortune SM, Rubin EJ. 2015. Phosphorylation of the Peptidoglycan Synthase PonA1 Governs the Rate of Polar Elongation in Mycobacteria. PLOS Pathogens 11:e1005010.

8. Kim J-H, Wei J-R, Wallach JB, Robbins RS, Rubin EJ, Schnappinger D. 2011. Protein inactivation in mycobacteria by controlled proteolysis and its application to deplete the beta subunit of RNA polymerase. Nucleic Acids Res 39:2210–2220.
